# Electromechanics of lipid-modulated gating of Kv channels

**DOI:** 10.1101/2020.06.12.051482

**Authors:** Nidhin Thomas, Kranthi K. Mandadapu, Ashutosh Agrawal

## Abstract

Experimental studies reveal that anionic lipid POPA and non-phospholipid cholesterol inhibit the gating of voltage-sensitive potassium (Kv) channels at 5–10% molar concentrations. Intriguingly, other anionic lipids similar to POPA, like POPG, have minimal impact on the gating of the same channels for reasons that remain obscure. Our long-timescale atomistic simulations show that POPA preferentially solvates the voltage sensor domains of Kv channels by direct electrostatic interactions between the positively charged arginine and negatively charged phosphate groups. Cholesterol solvates the voltage sensor domains through CH-*π* interactions between the cholesterol rings and the aromatic side chains of phenylalanine and tyrosine residues. A continuum electromechanical model predicts that POPA lipids may restrict the vertical motion of voltage-sensor domain through direct electrostatic interactions, while cholesterol may oppose the radial motion of the pore domain of the channel by increasing the mechanical rigidity of the membrane. The electromechanical model predictions are consistent with measurements of the activation curves of Kv channels for various lipids. The atomistic simulations also suggest that the solvation due to POPG is much weaker likely due to its bigger head-group size. Thus the channel activity appears to be tied to the local lipid environment, allowing lipids to regulate channel gating in low concentrations.

## Introduction

Voltage-gated Kv channels repolarize the neuronal membranes to restore the homeostatic potential across them and terminate the action potentials (1). The Kv channel is a tetrameric protein, which can be divided into two key domains based on their functionality (2–4) (Figure 1a). The first domain is the voltage sensor domain (VSD) that comprises of four segments: S1, S2, S3 and S4; see Figure 1a. The S4 segment carries multiple positively charged residues and is termed the voltage sensor as it responds to the changes in the electrochemical potential across the axonal membrane and initiates the opening of the channel. The S1, S2 and S3 segments carry the negative charges (also referred to as the counter charges), which regulate the movement of the S4 segment. The second key domain is the pore domain (PD) that comprises of S5, P helix (PH), and S6 segments (Figure 1a). These segments define the internal pore of the channel and form the passageway for ions to move across the membrane (5). According to the current working model, a change in the electrostatic potential across the membrane triggers an upward motion of the S4 segment, which in turn mobilizes the connected segments of the pore domain resulting in the opening of the channel (6, 7). While this traditional model is primarily protein-centric, a growing body of experimental studies now reveal that Kv channels show strong sensitivity to the lipid composition of the membranes which house the channels. A few selective lipids with contrasting properties have been recently shown to modulate the gating of Kv channels (8–12).

**Fig. 1.**
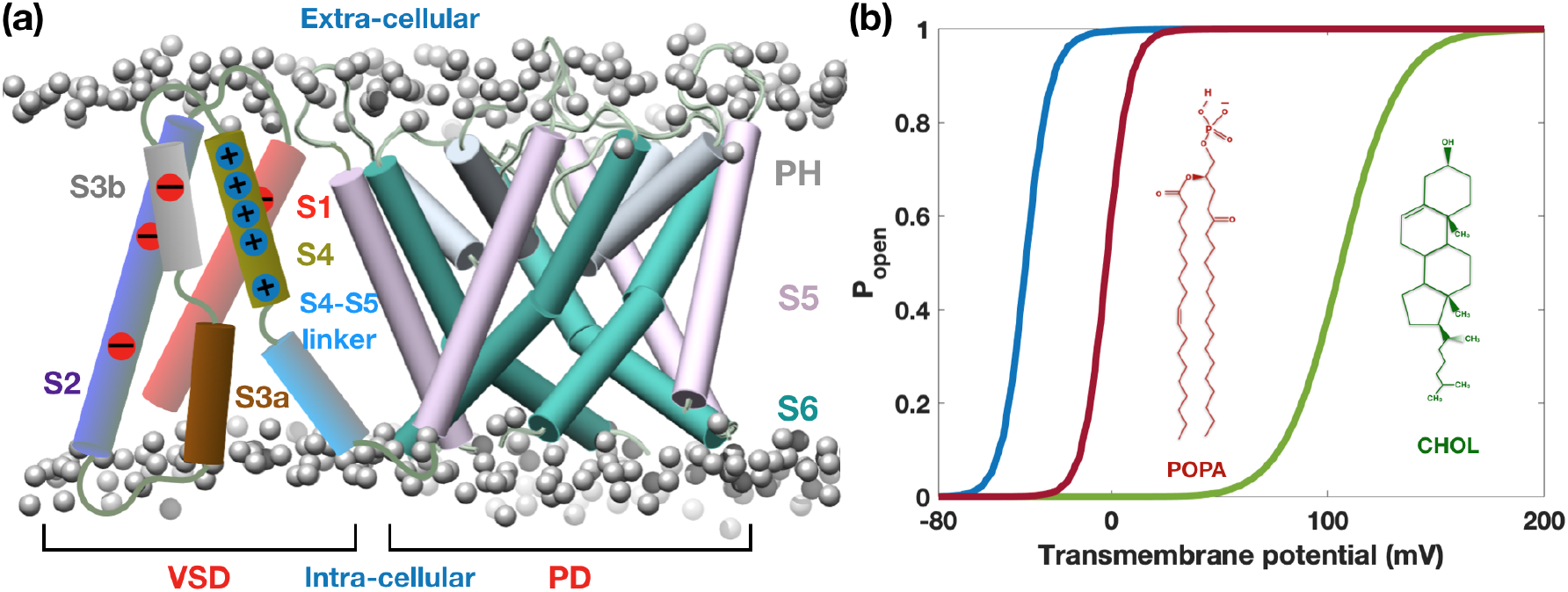
Lipid-dependent gating of Kv channels. (**a**) Schematic shows the key KvAP protein segments in the open configuration embedded in a lipid membrane. Segments S1-S2-S3-S4 constitute the voltage sensor domain (VSD) and segments S5-PH-S6 constitute the pore domain (PD). Segment S4, called the voltage sensor, contains the positively charged arginine (ARG) residues and responds to the changes in the transmembrane electrochemical potential triggering the opening of the channel. The subsequent increase in the pore size that allows the ions to flow across the membrane is regulated by the PD segments. (**b**) The gating response of the Kv channel is typically quantified by an activation curve (blue curve). In the resting state when the transmembrane potential is around −75 mV, the channel is in the closed state and the probability of the channel to be open *p_open_* ≈ 0. When the membrane depolarizes (negative potential decreases) and the activation voltage (voltage at which *p_open_* = 0.5) is crossed, the channel opens and *p_open_* ≈ 1. The presence of negatively charged lipid POPA and nonphospholipid cholesterol inhibits the opening of the channel and shifts the activation curves rightward (red and green curves show the qualitative trend). The mechanisms by which these specific lipids regulate channel gating is poorly understood. In this study, we systematically use all-atom molecular dynamics and continuum scale simulations to investigate this puzzle.

One of the key lipids that modulates the gating of Kv channels is the negatively charged phosphatidic acid (POPA) (10). A 25% concentration of POPA in DPhPC membrane is able to shift the activation voltage rightwards by ~30 mV, thus requiring larger depolarization to open the channels (10) (Figure 1b). What is remarkable is that even at a low concentration of 5%, POPA is able to generate a rightward shift of the activation voltage by ~10 mV (10). Furthermore, the impact of POPA on gating depends on its asymmetric distribution in the two leaflets (10). When present in the extracellular leaflet, POPA promotes opening of the channel, shifting the activation curve leftwards by ~15 mV. In contrast, when present in the intra-cellular leaflet, POPA shifts the activation curve rightwards by ~ 48mV, delaying the opening of the channel. Intriguingly, while POPA has significant impact, another lipid POPG with the same unit negative charge has much weaker influence on the gating of Kv channels (10). Similar to POPG, several other anionic lipids such as POPS, cardiolipin, PI and PIP with unit or more negative charges have a weaker impact on the gating of Kv channels compared to POPA (10).

Another important lipid that regulates the gating of Kv channels is cholesterol (CHOL), a non-phospholipid. Addition of CHOL in small percentages in DOPC membranes, arrests the gating of Kv channels. Addition of a mere 4% CHOL is enough to inhibit channel gating and shift the activation potential rightward by ~ 160 mV (9, 12); see Figure 1b. Furthermore, in stark contrast to POPA and CHOL, a positively charged lipid DOTAP, has also been shown to have a similar inhibitory effect on the gating of Kv gating (7, 8, 11).

The above experimental findings lead to several fundamental questions: why do POPA, CHOL and DOTAP selectively regulate the gating of Kv channels? Why do these specific lipids have a similar inhibitory effect? Do these lipids share any common physical property? Do these lipids invoke a common physical mechanism to regulate the channel gating? Lipid charge does not appear to be the common factor as POPA has a unit negative charge, DOTAP has a unit positive charge and cholesterol is charge neutral, and, yet, all of them inhibit the channel in a similar fashion. Moreover, there appears to be a lack of linear correlation between the head-group charge and the gating of Kv channel. This is evident from the fact that other anionic lipids with unit or higher negative charges, such as POPG, POPS and PIP lipids have much smaller impact than POPA lipids on the gating of Kv channels (10).

Another major mystery pertaining to these specific lipids is tied to the fact that they are able to regulate the channel gating at very low concentrations. It is not yet clear whether these lipids modify the overall behaviors of the membrane or do they alter the local properties in the vicinity of the channel? Formation of annular rings of cholesterol around the channel was hypothesized as a potential cause of cholesterol-induced inhibition of Kv channels (12). The validation of this hypothesis by either experiments or computations is till date pending. What is also not clear is whether these different lipids interact and arrest the motion of different protein segments in the ion channel or influence the same segment? It is conceivable that different lipids could potentially interact with distinct domains of the channel and influence the protein kinematics via distinct mechanisms.

In this work, we investigated the mechanisms of lipid-dependent voltage gating using a two pronged approach. First, we used long time scale all-atom molecular dynamics simulations to investigate the interactions of POPA, CHOL and POPG with the VSD of KvAP channel. Subsequently, we developed a continuum scale electromechanical model that accounts for electrostatic interactions and the elasticity of the membrane to quantify the impact of lipids on channel gating. Our studies reveal that both POPA and CHOL enter into the solvation shells of VSD protein segments and alter the local energetics of channel-membrane interactions. We find that POPA lipids aggregate in the vicinity of VSDs because of direct electrostatic interactions between the negatively charged phosphate group of POPA, and the positively charged arginine residues on S4. In contrast to POPA, we find that CHOL aggregates because of CH-*π* interactions between the CHOL rings and the aromatic rings of tyrosine (TYR) and phenylalanine (PHE) residues of the VSD segments. The electromechanical model suggests that POPA can regulate the vertical motion of S4 segment by direct electrostatic interactions while CHOL can make the bilayer locally stiff and restrict the opening of the pore domain. Despite having the same charge as POPA, POPG lipids show a much weaker propensity for solvating the VSD, likely because of a bigger head-group size. Coincidentally, both the solvating lipids, POPA and CHOL, have a small headgroup footprint, indicating that a small headgroup size might play a critical role in enabling lipids to solvate channel proteins. Overall, this general physical principle where lipids locally solvate and regulate conformational changes and functionality of proteins could be valuable in comprehending other vital lipid-protein interactions in cells.

## Insights from atomistic simulations

### POPA lipids solvate VSD because of ARG-phosphate electrostatic interactions

We began by studying the interactions between the VSD and the charged lipids using allatom molecular simulations. We used the crystallographic structure of the VSD of KvAP channel in the open configuration (13). We performed MD simulations of the VSD in a pure POPC bilayer and in a mixed bilayer comprising of 75% POPC lipids and 25% POPA lipids. Figure 2 shows a simulated system with the VSD embedded in the mixed bilayer (POPC lipids are shown in dark gray and POPA lipids are shown in white). The details of the MD simulations are provided in the Methods section and the SI.

**Fig. 2.**
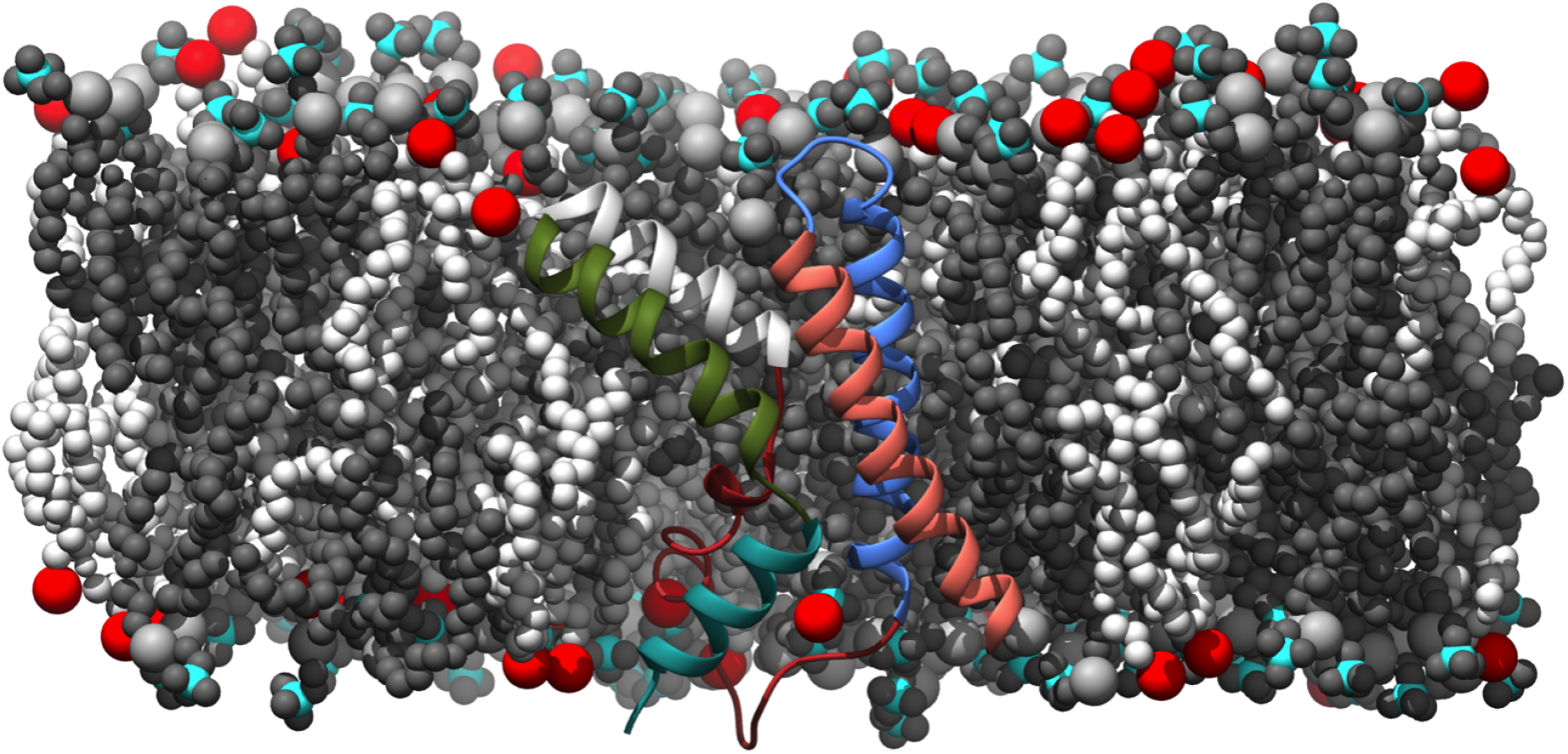
Molecular dynamics set-up used to study lipid-Kv channel interactions. Voltage sensor domain (VSD) of the KvAP channel is placed inside a POPC(75%)-POPA(25%) membrane. POPC and POPA lipids are colored gray and white, respectively. Nitrogen atoms of POPC lipids are colored cyan and the phosphorous atoms of POPA are shown in red.

We computed the two-dimensional radial distribution functions (RDFs), *g*_2*D*_(*r*), of POPC and POPA lipids around the VSD in the pure POPC and the POPC-POPA bilayers in order to quantify the local distribution of lipids around the VSD. The RDFs, in essence, quantify the areal density of lipids as a function of distance from the VSD (defined in Eqn. [1] in the Methods section). Figure 3a shows the RDFs of POPC lipids in pure POPC bilayer (gray curve), POPC lipids in POPC-POPA bilayer (cyan curve), and POPA lipids in POPC-POPA bilayer (red curve) with respect to the center of mass (COM) of the VSD. Compared to the POPC RDFs, the POPA RDF shows a peak within 1 nm. This demonstrates that POPA has a propensity to preferentially aggregate near the VSDs and enter the solvation shell within 1 nm of the protein. An inspection of the trajectory from our molecular simulations reveal that POPA lipids preferentially localize near the S4 segment, which contains the positive ARG residues.

**Fig. 3.**
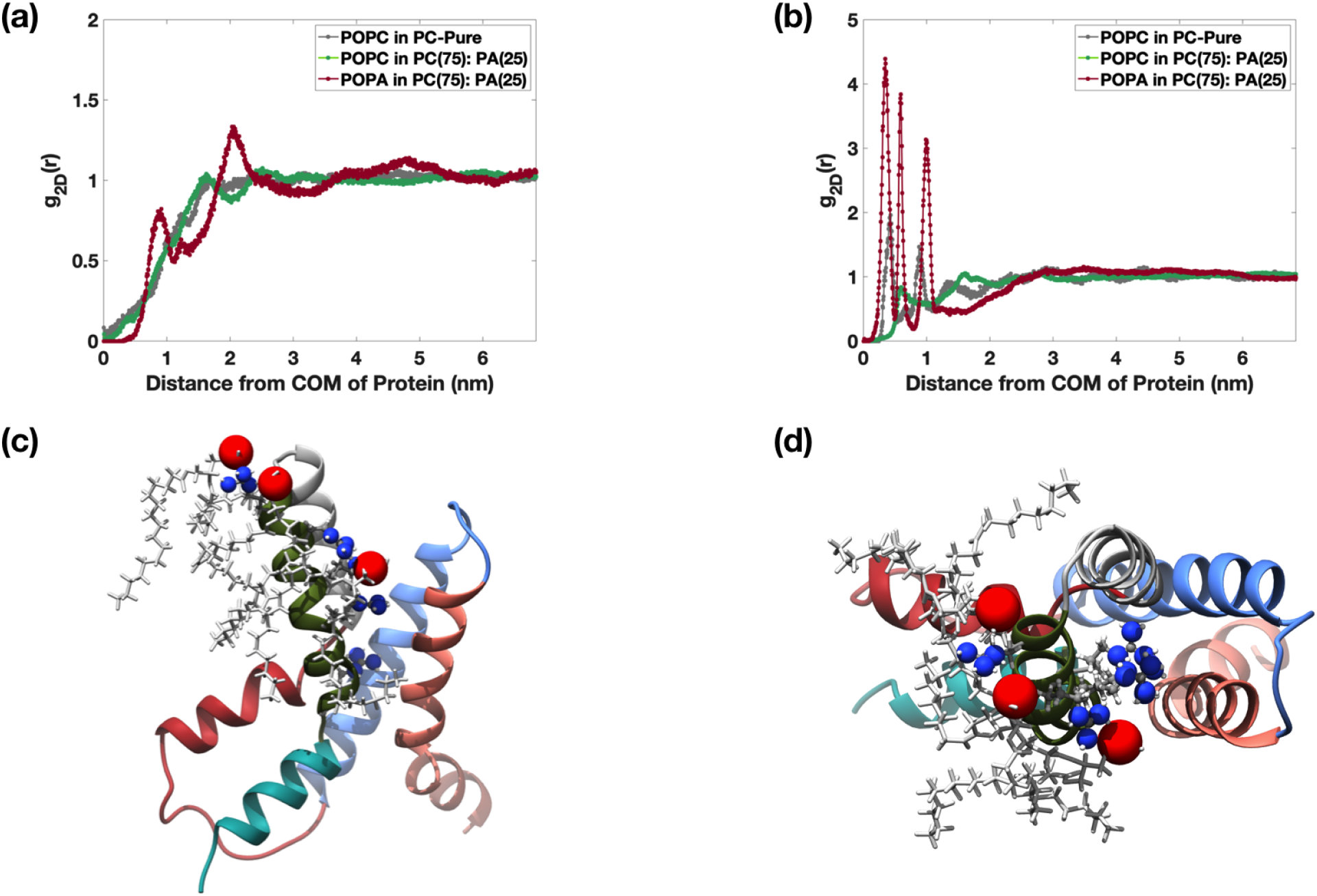
MD simulations reveal VSD solvation by POPA lipids (**a**) Radial distribution function (RDF) of POPA and POPC lipids around the centre of mass (COM) of VSD in the outer membrane leaflet. RDF of POPC lipids in a pure POPC membrane is in gray curve, RDFs of POPC and POPA in a mixed 75% POPC and 25% POPA membrane are in green and red curves, respectively. The peaks in the red curve reveal POPA aggregation and VSD solvation by POPA lipids. (**b**) RDFs of POPA and POPC lipids around the COM of the four S4 ARG residues (R117, R120, R123 and R126) in the outer leaflet of the pure POPC and mixed POPC-POPA membranes (same color coding as in (**a**)). The peaks in the red curve show aggregation of POPA lipids around the ARG residues in the S4 segment. (**c**) and (**d**) Side and top views showing clustered POPA lipids around the ARG residues in the outer leaflet. Phosphorous atoms of POPA and nitrogen atoms of ARG side chains are represented as red and blue spheres, respectively.

To quantify ARG-POPA interactions, we computed the RDFs of POPA and POPC lipids with respect to the ARG residues of the S4 helix as shown in Figure 3b. The POPA RDF with respect to ARG (same color coding) shows much higher peaks in the inner solvation shell compared to the RDF with respect to the COM of VSD in Figure 3a. Also, the POPA RDF has higher peaks compared to the POPC RDFs (same color coding) in Figure 3b. These plots indicate direct electrostatic interactions between the phosphate (P) groups of the POPA lipids and the ARG residues. Specifically, we find that the negatively charged phosphate group of the POPA lipids interact with the positively charged guanidinium group in the ARG side chain of the S4 helix. We also find that POPA aggregation is more predominant in the upper leaflet as the VSD is in the open configuration and the ARG residues are localized in the top leaflet of the membrane. The clustered POPA lipids around the ARG residues in the outer leaflet are shown in Figure 3c (side view) and Figure 3d (top view).

### Cholesterol solvates VSD because of CH-*π* interactions

Next, we investigated the interactions of CHOL with the VSD in a bilayer comprised of 75% POPC and 25% CHOL lipids. Figure 4a shows the RDFs of the CHOL and POPC lipids in the inner leaflet. The CHOL RDF shows a peak in the inner solvation shell revealing the propensity of CHOL to aggregate near the VSDs in the inner leaflet. The RDFs in the outer leaflet show weaker aggregation and are presented in Figure S1. This trend is in contrast to the POPA lipids which aggregate preferentially in the outer leaflet.

**Fig. 4.**
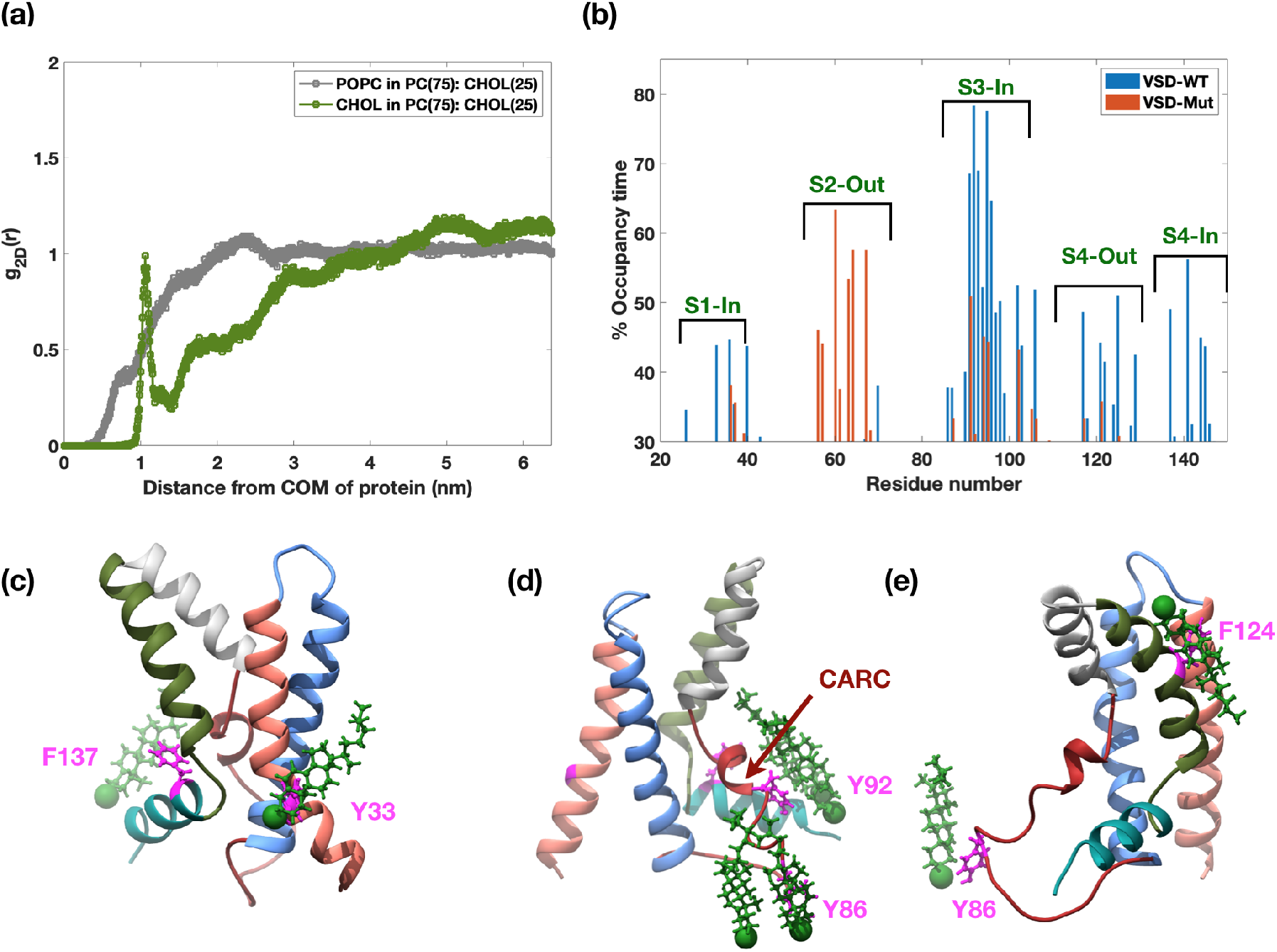
CHOL solvates VSD in the inner leaflet (**a**)) RDF of POPC (gray curve) and CHOL (green curve) around VSD in the inner leaflet of a mixed membrane with 75% POPC and 25% CHOL. The peak in the green curve shows preferential aggregation of CHOL around the VSD. (**b**) Occupancy time of CHOL around VSD for wild-type VSD and mutated VSD in which all aromatic side chains have been replaced with the alanine residue. The occupancy time is calculated as the percentage ratio of the total time each VSD residue spends within 6 angstrom of any CHOL molecule divided by the total sampling time. The blue histograms reveal the interaction of CHOL molecules with the aromatic side chains of PHE/TYR residues Y33 in S1 segment, Y86 and Y92 in S3 segment and F137 in S4 segment in the inner leaflet and F124 in S4 segment in the outer leaflet. Upon mutation, these interactions reduce significantly (red histograms). The only interaction in the mutated VSD occurs with R57 residue, which has a low binding affinity towards the polar head group of the CHOL molecule. (**c,d** and **e**) Three discrete simulation snapshots showing the interaction of CHOL (in green) with the aromatic rings of PHE and TYR residues (in magenta) and CARC domain (indicated by red arrow).

While POPA aggregation was dictated by electrostatic interactions between POPA and ARG residues, the cause of CHOL solvation is not very apparent. To investigate the specific interactions that dictate CHOL binding, we computed the percentage occupancy time of CHOL molecules with each VSD residue during the sampling time. We identified the residues within 6 angstroms of CHOL molecules during the sampling time using ‘gmx select’. We calculated the total time each residue spends within 6 angstrom of any CHOL molecule and divided by the total sampling time to compute the percentage occupancy time. Figure 4b shows the shortlisted residues which interact with the CHOL molecules for at least 30% of the sampling time. The figure shows that the CHOL molecules interact with the aromatic side chains of PHE/TYR residues Y33 in S1 segment, Y86 and Y92 in S3 segment and F137 in S4 segment in the inner leaflet and F124 in S4 segment in the outer leaflet. These CHOL interaction sites are marked with spikes (blue histograms) in Figure 4b.

To further validate the role of CH-*π* interactions in CHOL aggregation, we replaced all the aromatic side chains of the VSD with the alanine (ALA) residue and repeated the simulations. Upon mutation, CHOL occupancy time underwent a drastic reduction at the previously identified binding sites (red colored bars in Figure 4b). This confirmed the importance of CH-*π* interactions in regulating CHOL distribution. The only increase in occupancy time observed in the mutated state occurs around S2 segments. A closer examination reveals that S2 segment has a R57 residue, which has a low binding affinity towards the polar head group of the CHOL molecule. This interaction is likely the reason for CHOL aggregation in the mutant case. However, in the wild type structure, this interaction is probably overshadowed by the more powerful aromatic ring interactions. In addition to PHE or TYR residues, electrostatic interactions between the polar hydroxyl head group of CHOL molecules and the basic residues like ARG or lysine (LYS) could also facilitate CHOL binding. Such binding motifs, known as CRAC (Cholesterol Recognition Amino acid Motifs) and CARC (reverse order of CRAC sequence) domains, are known to promote interaction with the CHOL molecules (14–16). We identified one such CARC domain K88-Y92-L97 in the inner leaflet region of the S3 segment. The strong aggregation of CHOL around S3 in Figure 4b is likely because of a combined effect of the CARC domain and the CH-*π* interactions. The interactions of CHOL molecules with the PHE/TYR aromatic rings (in magenta) and the CARC domain (indicated by red arrow) at three discrete simulation times are shown in Figure 4c,d and e.

## Insights from the electromechanical continuum model

While atomistic studies reveal the preferential aggregation of POPA and CHOL around the VSD, the mechanisms by which they can potentially regulate channel gating remains unanswered. It is important to validate if the aggregated lipids can alter the energy landscape of the channel-membrane system so as to generate the experimentally observed shifts in the activation curves. We note that the shift of ~ 160 mV induced by CHOL is significantly larger than the shift of ~30 mV induced by POPA, suggesting that these lipids might be exploiting different mechanisms to regulate the gating. To gain such mechanistic insights and to quantify the consequences of solvation, it is necessary to transition to a continuum model due to the time and lengthscale limitations of the atomistic studies. However, quantifying the effect of lipids on the gating of a voltage-gated ion channel at the continuum scale is a unique problem as it requires a coupled electromechanical response of the channel-membrane system. On one hand, we have the evolving interactions of charges, and on the other hand, we have the mechanical motion of the pore-forming proteins leading to an elastic deformation of the neighboring membrane. While we have purely mechanical models of gating for a diverse set of channels in the literature (17–22), a combined electromechanical model of channel gating does not exist till date.

To predict the consequences of lipid solvation on channel gating, we develop a continuum scale electromechanical model that captures the essential channel-membrane interactions. We use the experimentally identified channel kinematics to judiciously construct a ‘model’ KvAP channel. Based on atomistic studies, we modify the membrane properties, and predict the consequences on channel configurations and the opening probability of the channel. Below are the main features and assumptions of the continuum model:

1. The ‘model’ channel is embedded in the membrane and accounts for the kinematics of the S4 segments in the VSDs and the S6 segments in the PD. Studies reveal that the key mechanical motion in a Kv channel is primarily associated with the S4 and S6 segments (6, 13, 23–26). The gating is initiated by the transmembrane S4 motion regulated by the electrostatic forces, followed by the radial motion of the S6 segments that increases the pore size allowing the ions to flow across the channel. Figure 5a shows the configurations of the S4 and S6 segments in the closed and open states of the channel. Using this discrete picture, we construct an equivalent ‘model’ channel shown in Figure 5b. Here, S4 is represented as a cylindrical protein (green) and the S6 segments are represented as a hollow cone (teal). Further details are discussed below and in the Methods section.
2. There are multiple competing mechanisms for the motion of the S4 domain outlined in the literature (6, 7, 27–31). Inspired from the commonality in these mechanisms, we assume that the representative S4 segment undergoes rigid body translation in the vertical direction and rotation about the vertical axis passing through the center of the cylinder (Figure 5b). The S4 segment configuration in turn regulates the positions of the ARG residues present on it. As a consequence, movement of S4 regulates the salt-bridge connections ARG residues make with the counter charges present on the S1, S2 and S3 segments (see Figure S2 and Table S1). We assume that the counter charges remain stationary during the gating movement based on the proposed configurations of the S1, S2, and S3b segments in the open and closed states (6, 13, 23, 24, 32) (see SI for additional details).
3. The PD model is based on the known structure and kinematics of the pore-forming S6 segments (23–26, 33). The PD model is assumed to be made of two domains: one in the outer leaflet of the membrane and the other in the inner leaflet of the membrane (Figure 5b). Based on the experimental findings (6, 25), we assume that the outer leaflet domain is a hollow cone that remains stationary during the gating of the channel. The bottom leaflet domain is also a hollow cone but it undergoes configurational changes as the channel transitions between the open and closed states. While the closed structure of the Kv channels is not yet established, the current hypothesized model in the literature assumes that the bottom cone becomes more conical, which increases the vertical height and reduces the pore size of the PD in the closed configuration (2, 23, 24, 26, 34, 35). We model this effective change in the geometry of the bottom domain as shown in Figure 5b (see Figure S3 for more details). Lastly, we assume that the motion of this bottom domain is rigidly coupled to the motion of the S4 segment via a deformation mapping.
4. As the PD transitions between the open and closed states, the surrounding lipids reconfigure to maintain proximity to the PD and avoid exposure to the surrounding water. Thus, the gating movement of the PD is associated with an energetic cost to remodel the neighboring membrane. To quantify this contribution, we model lipid membrane as a 2D elastic sheet. The midplane of the membrane is assumed to remain flat and the two leaflets are assumed to be kinematically decoupled. The geometry of the membrane at the protein interface is assumed to be regulated by the PD geometry. Since the top domain of the PD has a fixed conical geometry, the outer leaflet of the membrane does not undergo any remodeling during channel gating (see Figure 5b). In contrast, the bottom domain of the PD perturbs the orientation and the thickness of the inner leaflet as it transitions between the closed and open states (see Figure 5b).
5. The free energy of the system, based on the discussion above, comprises of the electrostatic energy and the mechanical energy. The electrostatic contributions arise from the transmembrane potential energy, direct electrostatic interactions between the ARGs and the counter charges and the ARGs and the anionic lipids, and self energy of the charges (Eqns. [2]–[8] in the Methods section). The mechanical energy contribution comes from bending and thinning of the membrane (Eqns. [9]–[10] in the Methods section). The details of the model are discussed in the Methods section and the SI.

**Fig. 5.**
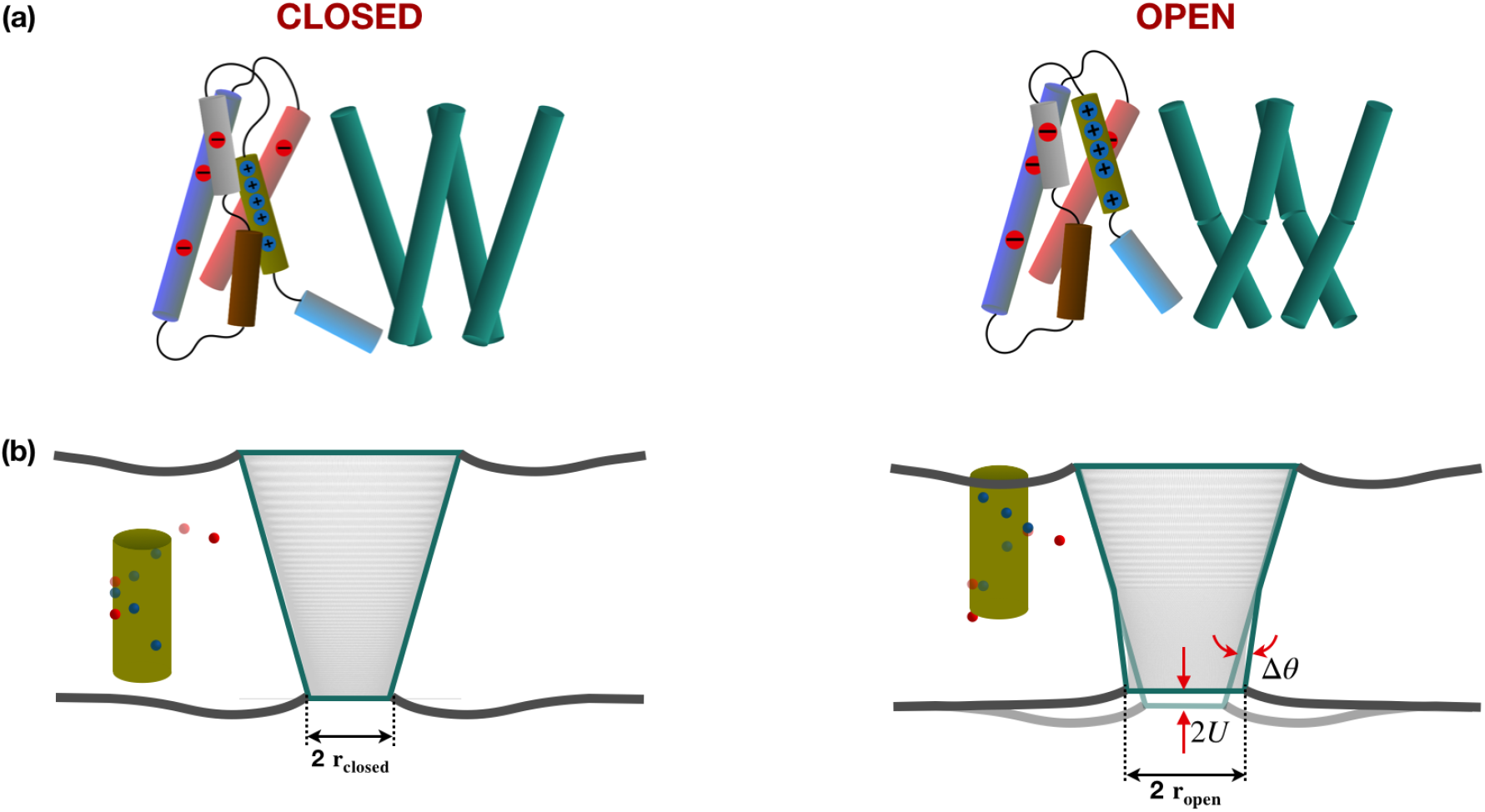
Schematic of the electromechanical ‘model’ of Kv channel. (**a**) Configuration of the helical segments of the VSD and the PD (selective segments shown) in the closed and open states used to set-up the ‘model’ Kv channel (motivated by findings in (23–26)). In the closed state, the S4 segment is closer to the inner leaflet and the PD helices are tilted with respect to the vertical axis. During opening of the channel, S4 helix rotates and moves up within the membrane. All other helices in the VSD essentially remain stationary. Concurrently, the C-terminal segment of the S6 helix tilts and twists to open the channel. (**b**) Schematic of the ‘model’ Kv channel employed in this study to predict lipid-dependent gating. The model retains the ARG residues (blue circles) and the counter charges (red circles) in the VSD and a hollow conical PD. The left and right panels show the closed and open configurations, respectively. The vertical translation of ARGs changes the electrostatic interactions with the counter charges and the charged lipids. The tilting of the PD changes the lipid orientation (Δ*θ*) and membrane thickness (2*U*) at the protein interface. The associated electrostatic and mechanical energies regulate the gating of the channel.

### POPA can regulate the vertical translation of S4 by direct electrostatic interactions

With the ‘model’ ion channel, we first predicted the consequences of channel solvation by POPA lipids. We model POPA’s effect via direct electrostatic interactions with the ARG residues. We do not invoke any POPA-induced changes in the mechanical properties of the membrane as we are not aware of any such findings in the literature. For the POPA-solvated state, we placed one POPA lipid within 1 nm and the other lipid within 1.5 nm from the surface of S4 helix in the outer leaflet of the membrane based on the RDF presented in Figure 3a (see Table S2). Since the closed configuration of the VSD is unavailable, we placed two POPA lipids at the similar distance from the S4 in the inner leaflet as in the outer leaflet. In addition, we placed an extra POPA lipid within 1 nm motivated by the fact that the RDF of POPA around the LYS residue in the inner leaflet shows POPA aggregation (Figure S4).

Figure 6 shows the impact of the POPA lipids on the gating of the ‘model’ KvAP channel. Figure 6a and Figure 6b show the total energy landscapes for the model KvAP channel in the pure POPC and mixed POPC-POPA membranes, respectively, at 0 mV transmembrane potential as a function of the vertical translation (x-axis) and the azimuthal rotation (y-axis) of the S4 segment. The predicted energy values at the energy wells in *k_B_T* are shown in white text. The rightmost energy well marked by an arrow corresponds to the open state of the channel. All the remaining energy wells correspond to the closed state. We compute the energy landscapes at different transmembrane potentials and estimate the opening probability curves (defined in Eqn. [11] in the Methods section; also referred to as activation curves) for the KvAP channel in pure POPC and POPC-POPA bilayers. Figure 6c shows the activation curves for the two systems (pure POPC in gray curve; POPC-POPA in red curve). The activation curves for the POPC-POPA bilayer show a rightward shift compared to the pure POPC bilayer curves (red curve in Figure 6c). The activation voltage (voltage at which the opening probability becomes half) for the pure POPC and POPC-POPA bilayers are −30 mV and 1 mV. Thus, POPA solvation leads to a rightward shift of 31 mV.

**Fig. 6.**
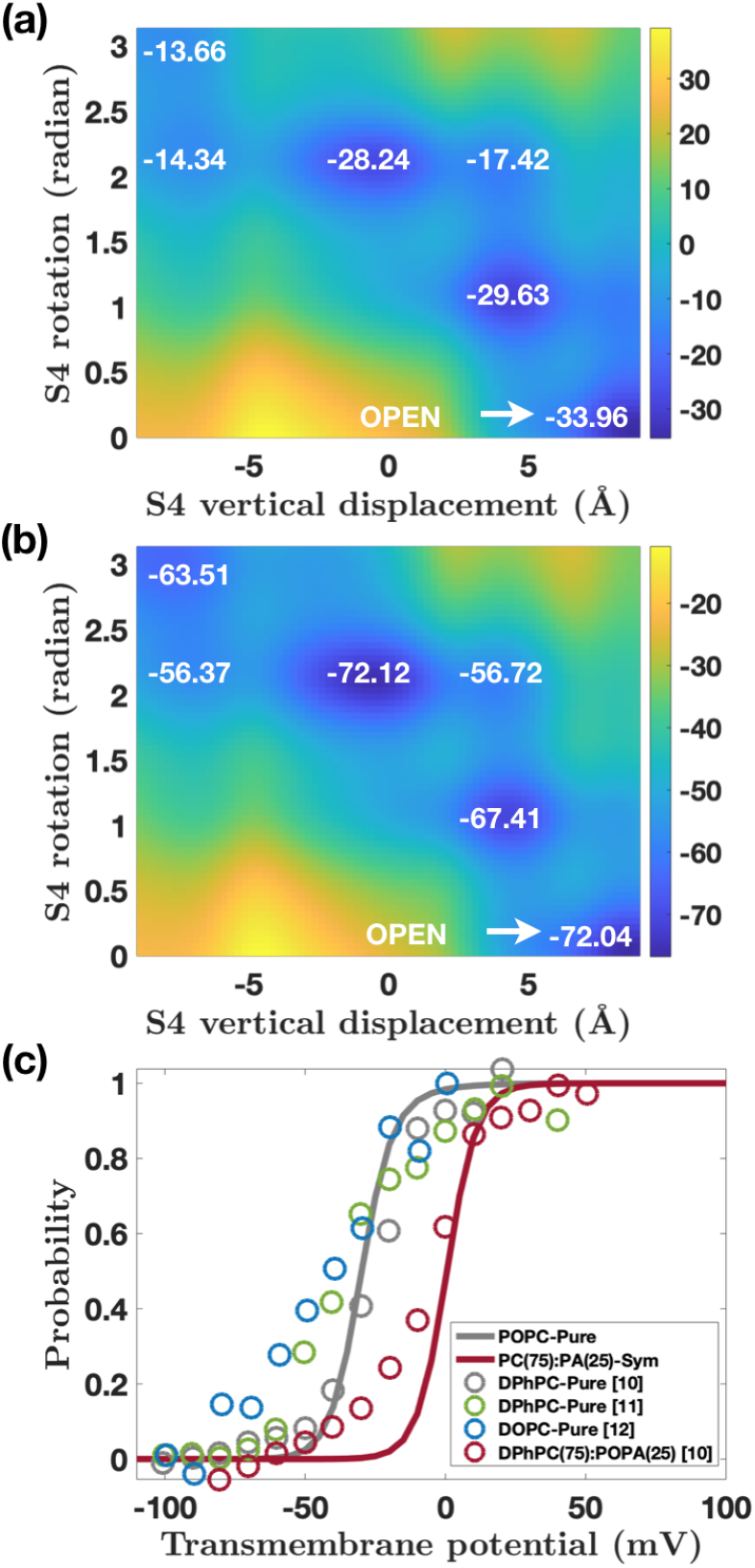
POPA regulates the gating of ‘model’ Kv channels by direct electrostatic interactions. (**a** and **b**) Energy landscapes of the KvAP channel at 0 mV transmembrane potential in pure POPC and mixed POPC-POPA membranes. The vertical position of S4 is on the x-axis, the rotation of S4 is on the y-axis, and the energy values (in *k_B_T*) at the wells are in white text. The rightmost well correspond to the open state of the channel. All other energy wells correspond to the closed states. POPA-ARG interactions lower the energies of the wells in (**b**). (**c**) Predicted activation curves for the KvAP channel in pure POPC membrane (gray curve) and POPC-POPA membrane (red curve). Presence of POPA creates a rightward shift of 31 mV. Experimental data for KvAP channels in DPhPC membranes (gray and green circles), DOPC membranes (blue circles), and DPhPC-POPA membranes (red circles) show good agreement with the model predictions. Experimental data from *eLife* (10); *Journal of Biological Chemistry* (11); and *Advances in Experimental Medicine and Biology* (12).

To understand these trends, we can analyze the energy landscapes in Figure 6a and Figure 6b. For the pure POPC bilayer, the open state well has the least energy of −33.96*k_B_T*. As a result, the channel prefers to be in the open state at 0 mV. When POPA solvates the VSD, the favorable electrostatic interaction between the POPA and ARG residues in the two leaflets reduce the energies of the wells. The energy of the open state well changes to −72.04 *k_B_T*, resulting in a change of −38.08 *k_B_T*. But in comparison, the energies of the closed state wells undergo a larger reduction (ranging from −37.78 *k_B_T* to −49.85 *k_B_T*) as the POPA solvation is stronger in the inner leaflet. So, even though the open state continues to have the least energy, the closed state wells become energetically more favorable. As a result, the channel develops a higher propensity to be in the closed states at 0 mV. This reduced probability of the channel to be in the open state manifests itself in the form of the rightward shift in the activation curve.

The experimental data for the KvAP channel are available for the DOPC, DPhPC, and DPhPC-POPA membranes (8, 10–12). While we do not have exact data for the POPC and POPC-POPA membranes, we can expect the channel response to be similar in POPC, DOPC, and DPhPC membranes because of two reasons. First, POPC membrane has an area compressibility modulus of 255 mN/m, which is comparable to the compressibility modulus of 265 mN/m of the DOPC membrane (36–38). Second, when we plot the experimental data for the DOPC and DPhPC membranes for the KvAP channel, they show significant overlap. This indicates that the membrane properties are comparable and the channel operates in a similar manner in the two membranes. Motivated by these arguments, we overlaid the available experimental data on to our predicted curves. The experimental data for the KvAP channel in DOPC membrane (blue circles) (12), DPhPC membranes (gray and green circles) (10, 11) and DPhPC-POPA membrane (red circles) (10) are presented in Figure 6c. Model predictions are in excellent agreement with the experimental data. This suggests that the direct electrostatic interactions between the POPA and the ARG residues could be a potential mechanism by which POPA regulates gating of Kv channels.

### CHOL can regulate the radial movement of PD by local stiffening of membrane

Next, we estimated the consequences of channel solvation by CHOL. Studies have shown that CHOL stiffens POPC/DOPC membranes between 2-3 times (39–42). Hence, we increased the bending modulus 2 times and the compression modulus 2.5 times (Table S3) to account for the effect of CHOL solvation. Modeling the change in the elasticity of the membrane, we computed the energy landscapes and the activation curves for the KvAP channel in the presence of CHOL.

Figure 7a and Figure 7b show the energy landscapes in the pure POPC and POPC-CHOL membranes, respectively. The open state well has the least energy for the pure POPC membrane system. In the presence of CHOL, the energy of the open state well goes up making the closed states energetically favorable. This is so because the open state of the channel is associated with a higher compression of the inner leaflet in the protein vicinity. Hence, CHOL-rich membrane with increased stiffness, opposes thinning and makes the open state energetically costlier. As a consequence, the channel develops a higher propensity to be in the closed states. This results in a rightward shift of 155 mV in the activation curve shown in Figure 7c (pure POPC in gray curve and POPC-CHOL in green curve). This prediction is well aligned with the experimentally measured shift of 160 mV for the KvAP channel induced by addition of ~ 4.0% molar concentration of CHOL in DOPC membrane (pure DOPC in gray circles and DOPC-CHOL in green circles) (12). The agreement between the model predictions and the experimental data suggests that the CHOL-induced changes in the elasticity of the membrane could be a key factor regulating channel conformations.

**Fig. 7.**
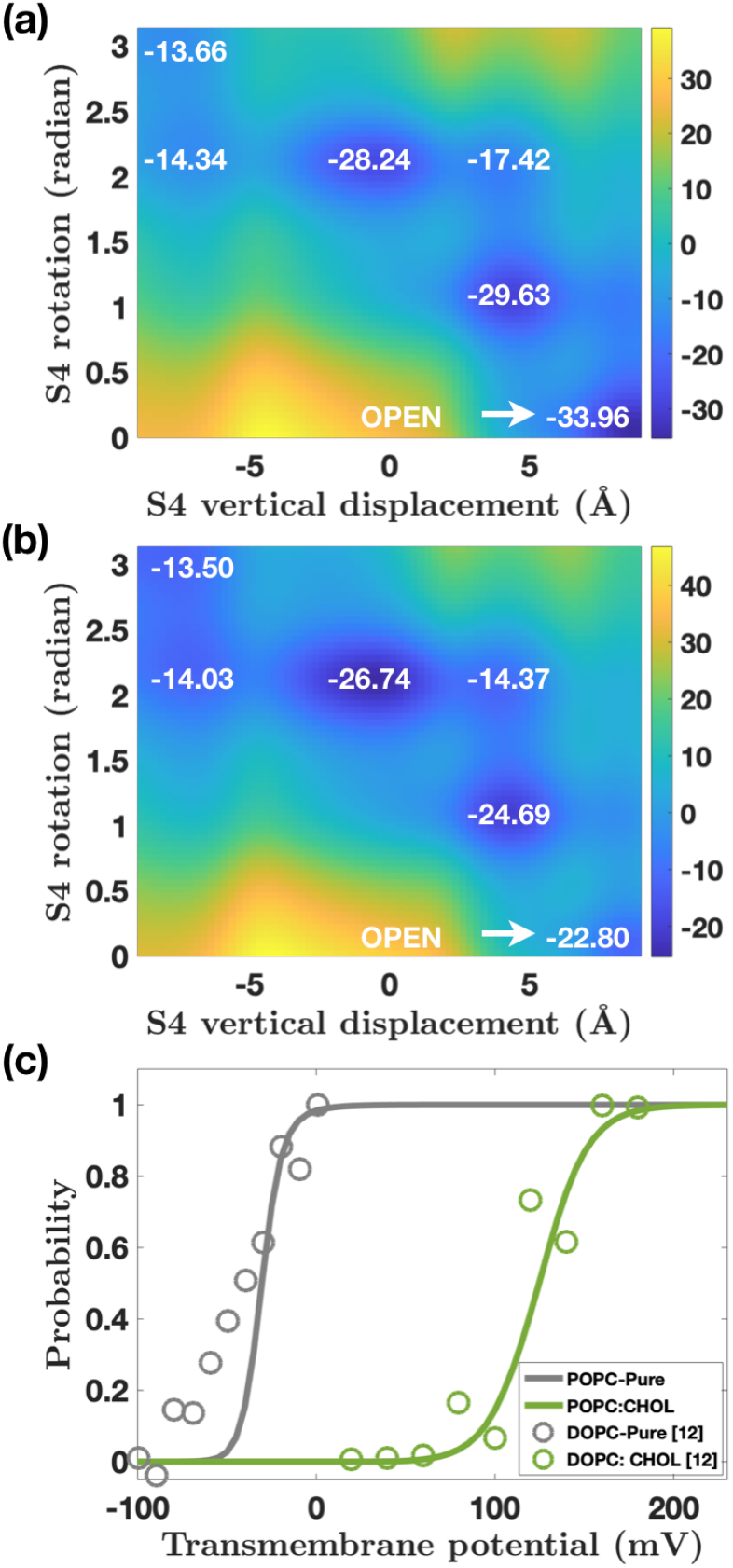
CHOL regulates ‘model’ Kv channel gating by restricting PD motion. (**a** and **b**) Energy landscapes of the KvAP channel at 0 mV transmembrane potential in pure POPC and mixed POPC-CHOL membranes. The vertical position of S4 is on the x-axis, the rotation of S4 is on the y-axis, and the energy values (in *k_B_T*) at the wells are in white text. The effect of CHOL was modeled via increased bending and compression moduli for the mixed POPC-CHOL membrane. CHOL penalizes the open state energy well in (**b**) due to increased membrane deformation associated with the open state of the channel. (**c**) Predicted activation curves for the KvAP channel in POPC membrane (gray curve) and POPC-CHOL membrane (green curve). Presence of CHOL creates a rightward shift of 155 mV. Experimental data for the KvAP channels in DOPC membranes and DOPC-CHOL membranes are shown in gray and green circles, respectively. The experimental data and the modeling predictions show good agreement. Note. Experimental data from *Advances in Experimental Medicine and Biology* (12).

## Discussion

Phospholipids are essential for the proper gating dynamics of voltage-gated Kv channels. Low concentrations of POPA and CHOL have been shown to inhibit Kv channel gating. Here, we used molecular dynamics simulations and a continuum electromechanical model to identify the biophysical mechanisms. Our atomistic study reveals that POPA solvates the VSD because of negative charge, and CHOL solvates the VSD because of CH-*π* interactions. Our continuum analysis suggests that solvated POPA can regulate the motion of S4 segment via direct electrostatic interactions, and solvated CHOL can regulate the motion of the pore domain by changing membrane elasticity. Overall, our findings suggest that the channel gating is regulated by lipids that are able to locally solvate the channel and alter the energetic landscape of the channel-membrane system.

### POPG lipids show weaker solvation

Atomistic study shows that POPA lipids preferentially aggregate near VSDs because of direct electrostatic interactions between the negative charged headgroup and the positively charged ARG residues. However, if the charge interactions were the sole reason for solvation, any lipid with a negative charge should be able to solvate the VSD. To test this conjecture, we repeated our analysis for a mixed bilayer comprising of 75% POPC and 25% POPG lipids. The RDFs for the POPG (blue curve) and POPC (cyan curve) lipids with respect to the COM of the VSD and the ARG residues are shown in Figure 8a and Figure 8b, respectively. The RDFs of the POPA lipids from Figure 3 are reproduced to present a comparative picture. The two sets of plots in Figure 8 reveal that unlike POPA RDF, POPG RDF does not exhibit peaks in the inner solvation shell. This suggests that despite the same charge, POPG lipids exhibit much weaker electrostatic interactions with the ARG residues, which results in a weaker solvation of the VSD.

**Fig. 8.**
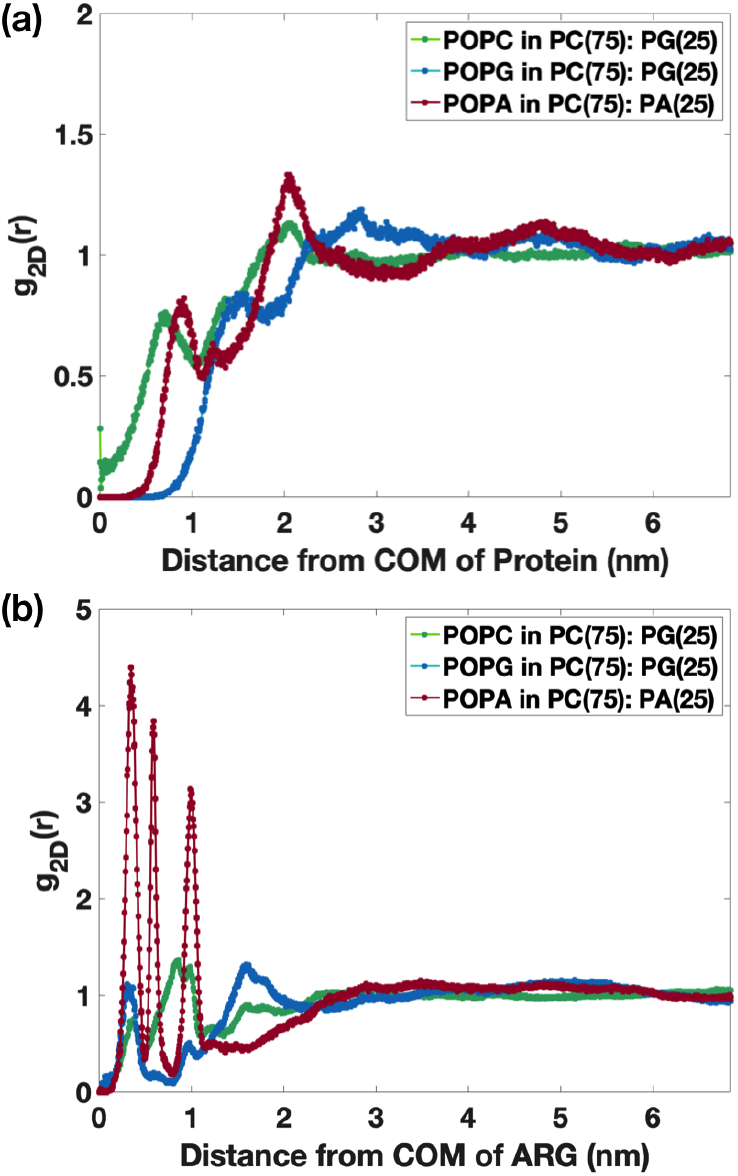
POPG lipids show weaker aggregation around VSD compared to POPA lipids despite having the same unit negative charge. (**a**) RDFs of POPG lipids (blue curve) and POPC lipids (green curve) around the COM of VSD in the outer leaflet of a membrane comprised of 75% POPC and 25% POPG lipids. RDF of POPA (red curve) is reproduced from Figure 3a. A comparison of the red and blue curves shows that POPA lipids exhibit closer and stronger aggregation around VSD than POPG lipids. (**b**) RDFs of POPG and POPC lipids around the COM of the four S4 ARG residues (R117, R120, R123 and R126) in the outer leaflet of the POPC-POPG membrane (same color coding as in (**a**)). RDF of POPA (red curve) is reproduced from Figure 3b. The peaks in the red curve are much stronger than the peaks of the blue curve showing preferential aggregation of POPA lipids around the ARG residues compared to POPG lipids.

We can analyze the structural properties of POPA and POPG to gain mechanistic insights into the different solvation levels they exhibit. A comparison of the headgroup sizes of POPA and POPG lipids shows that POPA lipids have a 35% smaller volume than POPG lipids (see Figure S5) (43–46). While POPA has a hydrogen ion linked to the phosphate group, POPG has a glycerol group connected to the phosphate group, resulting in a bigger headgroup and a lower charge density. As such, we hypothesized that the POPG lipids should exhibit smaller residence time compared to POPA lipids in the vicinity of the ARG residues. We computed the total residence time of POPC, POPA and POPG lipids within 0.5 nm of the COM of the VSD ARG residues (Figure S6). While POPA lipids have a residence time of upto 1500 ns (Figure S6a), POPG lipids have a residence time of only upto 300 ns (Figure S6b). As POPA and POPG lipids have identical acyl chains, this marked reduction in the residence time of POPG lipids can potentially be attributed to the reduced electrostatic interaction resulting from a bigger headgroup size. This analysis, therefore, suggests that the headgroup size may be a key parameter that determines the strength of lipid-protein interactions and the extent of protein solvation by lipids.

### Cholesterol can solvate pore domain because of aromatic rings and CARC/CRAC domains

Our molecular simulations show that the VSD possesses aromatic rings and CARC domain that allows CHOL to aggregate in the channel vicinity. We further investigated the crystallographic structure of the KvAP PD (2) to identify additional CHOL binding sites. We found that the PD contains a much larger number of CHOL binding sites than the VSD. Figure 9 shows all the aromatic rings present in the PD. These rings are present in i) the PHE residues (red color), ii) the TYR residues (blue color), and iii) the tryptophan (TRP) residues (tan color). While the aromatic rings in the PHE/TYR residues are accessible to CHOL molecules, the rings in the TRP residues are present close to the selectivity filter and hence, are not amenable to CHOL binding. In addition, PD contains three CARC domains and one CRAC domain, three of which are present in the inner leaflet region (see Table S4). The presence of these sites strongly suggests that CHOL is likely to exhibit stronger solvation in the presence of the entire structure of the Kv channels.

**Fig. 9.**
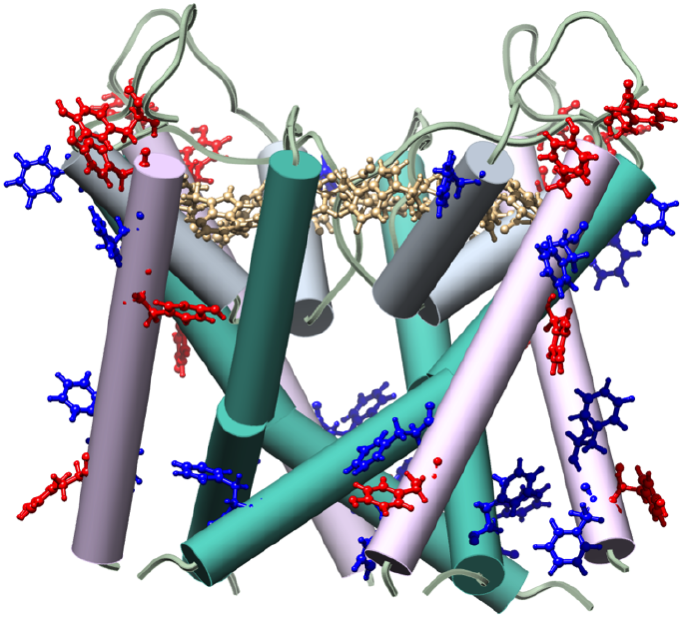
Aromatic side chains in the PD of the KvAP channel. Aromatic side chains in PHE residues (red color) and TYR residues (blue color) can interact with the CHOL molecules. The aromatic side chains in TRP residues (in tan color) close to the selectivity filter are likely to have minimal interactions with CHOL because of steric hindrance.

### Electromechanical model predicts DOTAP-mediated inhibition of KvAP gating

In addition to POPA and CHOL, positively charged lipid DOTAP has been shown to inhibit KvAP gating (7, 8, 11). MD simulations reveal that DOTAP lipids move away from the S4 helix because of the repulsive interactions between the positive charges on the DOTAP and the S4 segment (47). Furthermore, higher water penetration was observed in the S4 vicinity (47). We employed our electromechanical continuum model to quantify the consequences of these findings on channel gating. Since DOTAP leads to increased water penetration, we increased the dielectric constant of the lipid-protein interface region in the model. This led to a reduction in the electrostatic interactions between the S4 ARG residues and the counter charges. Since the mechanical energy favors the closed states of the channel, a reduction in the electrostatic energy made the closed states relatively energetically more favorable. As a result, a larger depolarization is required to open the channel. This results in a rightward shift in the activation curve (Figure 10). Figure S7a and Figure S7b show the energy landscapes for the pure POPC and POPC-DOTAP membranes for the KvAP channel. Invoking a dielectric constant of 20 (compared to 14.5, used in the prior calculations), the activation curves predicted by the model (pure POPC in gray curve and POPC-DOTAP in red curve) match very closely with the measured experimental curves for KvAP channels in DPhPC and POPC-PG-DOTAP membranes (pure DPhPC in gray circles, DPhPC-DOTAP in red circles, POPC-PG-DOTAP in green circles) (8, 11). The model predicts a rightward shift of 44 mV, which is close to the 38 mV shift observed in experiments (11). By invoking a dielectric constant of 30, the predicted model curve (purple curve) matches well with the experimental curve obtained in POPE-POPG-DOTAP membranes (purple circles) (7). The predicted rightward shift of 110 mV matches well with the experimentally observed shift of 107.5 mV.

**Fig. 10.**
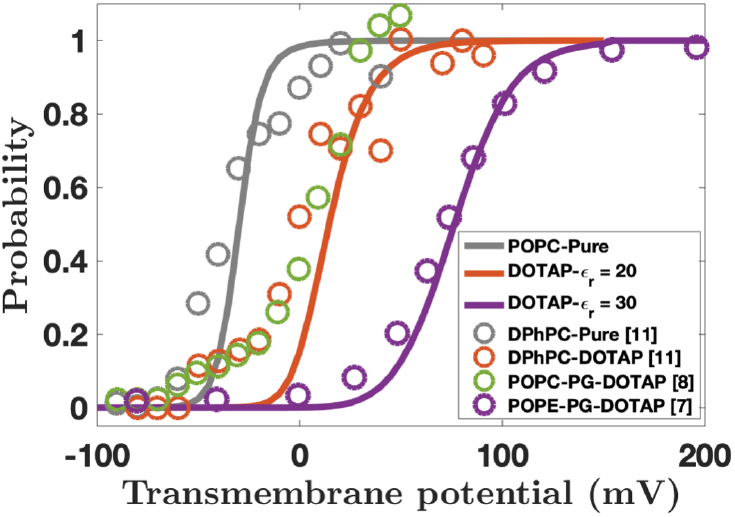
Predicted activation curves for the KvAP channel in pure POPC membrane (gray curve) and POPC-DOTAP membranes (red and purple curves). The red and purple curves correspond to dielectric constants of 20 and 30, respectively. Experimental data for DPhPC-DOTAP membranes, POPC-PG-DOTAP membranes, and POPE-POPG-DOTAP membranes are shown in red, green and purple circles, respectively. Note. Experimental data from *Journal of Biological Chemistry*(11); *Nature* (8); and *Nature Structural & Molecular Biology*(7).

### Limitations

While computational studies continue to enrich our mechanistic insight into the working of Kv channels, they are limited by simplifications and subject to experimental validation (48). For example, in this study, one of the key limitations is the lack of MD simulations for the closed configuration of the Kv channels. While we can make cautious extrapolations about lipid-protein interactions in the closed configuration based on the simulations performed on the open configuration of the VSD, performing explicit simulations with the closed configuration, when the crystal structure becomes available, would definitely add to our insights. The second limitation pertains to the continuum model. The channel is a complex protein structure that undergoes numerous degrees of motion and interacts with lipids in multiple ways. Here, we have restricted ourselves to a minimal set of kinematics that captures the essence of the lipid-protein interactions. In the future, it would be desirable to refine the model and incorporate more degrees of freedom such as twisting and tilting of lipids around the protein segments.

Also, the cryo-EM structure of the KvAP channel published recently (49) shows that KvAP channel is a non-domain swapped channel in which the S6 helix in the PD does not have a pronounced kink in the open configuration. The kinematics invoked in our electromechanical model is based on the domain-swapped structure of the KvAP channel discovered earlier in which the S6 helix was kinked near G220 residue in the open configuration (26). It would be insightful to adjust the kinematic constraints and revisit the model predictions in the light of the new findings in the future studies.

### General principles

While we investigated the interactions of POPA and CHOL with KvAP channel in this study, similar lipid-channel interactions have been found to be critical for the working of other types of channels. For example, POPA lipids have been shown to inhibit the gating of Kv Chimera channels (10). POPA lipids have also been found to stabilize the tetrameric assembly of KcsA channels (50–52) and show stronger interaction with KcSA channels than POPG lipids (50, 53, 54). Similarly, CHOL has also been shown to inhibit the gating of Kv1.3 and Kir2.2 channels (16, 55–57). Going beyond Kv channels, POPA lipids have been shown to inhibit the gating of epithelial Na channels (58). Furthermore, VSDs of Na channel homologs have been shown to possess high sensitivity towards anionic lipids (59). Since the findings in this study reveal general physical principles by which lipids could interact and modulate ion channels, our study could be of value to gaining mechanistic insights into the working of these other types of channels.

## Methods and Materials

### Molecular Dynamics Simulations

We used CHARMM-GUI (60, 61) to create an all-atom model of initial configurations. The crystallographic structure of the voltage sensor domain (VSD) of the KvAP channel in open configuration was obtained from (13). Five systems were simulated to study the effect of anionic lipids and cholesterol (CHOL) in KvAP channel (see Table S5). In the first system, VSD was inserted into pure POPC lipid bilayer. 601 POPC lipids were used to create the bilayer. 300 POPC lipids were used to create the top leaflet and 301 lipids were used in the bottom leaflet. The additional lipids were used to balance the area per lipid of each leaflet in order to maintain zero surface tension in each leaflet. The position of the center of mass (COM) of VSD with respect to the center of bilayer was compared with the results provided by (13). Other systems were created in CHARMM-GUI by replacing 25% of POPC with anionic lipids (POPA and POPG) or CHOL. We used CHARMM36m force field (62, 63) to run simulations in GROMACS 2018.3 (64).

The radial distribution function (RDF) *g_2D_* (**r**) represents the two-dimensional distribution of phosphorus atoms around either the COM of the VSD or the ARG residues. The RDFs were calculated using ‘gmx rdf’ command in the GROMACS. We used the center of mass (COM) of the entire VSD and the COM of the ARG group as the reference group to compute RDFs. We used the phosphorus atoms of individual lipids as the selection groups. For RDFs between the ARG residues and the phosphate groups, COM of the first four ARG residues (R117, R120, R123, R126) and the phosphorus atoms in the top leaflet were used. The RDF *g*_2*D*_(**r**) is defined by

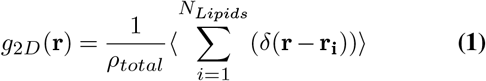

where 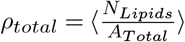 in which *N_Lipids_* is the total number of lipids in the bilayer and *A_Total_* is the surface area of the membrane patch. Above, **r** is the position vector where *g*_2*D*_ is evaluated, **r**_*i*_ is the position of the *i^th^* phosphorus atom with respect to the COM of the protein, and 〈.〉 is the sampling average. We compute the RDFs as a function of time to ensure that RDFs have converged. Two sample RDFs for POPC(75):POPA(25) and POPC(75):POPG(25) systems are shown in Figure S8. The fluctuations in the RDFs fall below 10% at the end of 2 *μs*, indicating convergence of RDFs.

### Energy contributions in the electromechanical model

The free energy of the total membrane-channel system is given by

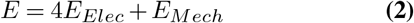

where *E_Elec_* is the electrostatic energy of a single VSD and *E_Mech_* is the mechanical energy of the surrounding membrane. A factor of 4 in front of *E_Elec_* is incorporated to account for the 4 VSD subunits. The electrostatic energy *E_elec_* is composed of four components

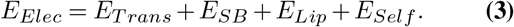

We discuss these contributions in greater detail next.

#### Transmembrane potential energy (*E_Trans_*)

The electrostatic energy due to the transmembrane potential arises from the changing vertical positions of the ARG residues in a uniform external electric field acting across the membrane. We assume the potential at the interface of the outer leaflet with water to be at 0 mV and vary the potential at the interface of the inner leaflet with water from −150 mV to 200 mV. Since the counter charges are fixed, their contribution does not change with the S4 movement and hence is suppressed in the model. We thus write the electrostatic energy due to the transmembrane potential as

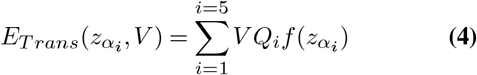

where *f*(*z_α_i__*) is given by

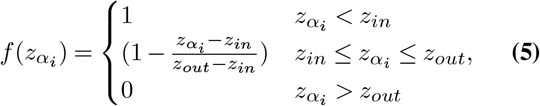

*V* is the transmembrane potential across the membrane, *Q_i_* is the charge of the *i_th_* ARG, *z_α_i__* is the vertical position of the *i_th_* ARG measured from the center of the bilayer (Eqn. [S3]), and *z_in_* = −16 Å and *z_out_* = 16 Å define the height of the interfaces up till which water is able to penetrate into the two leaflets.

#### Salt bridge energy (*E_SB_*)

The salt bridges ARG residues form in the open and the closed states for the KvAP channel are listed in Table S1. There are three major salt bridge connections in the open configuration (E45-R126, D62-R133 and E107-R123) and the closed configuration (E45-R117,D62-R120, and D72-R126) for the KvAP channel (Figure S11). The salt bridge interactions in the open configuration were obtained from (13). The salt bridge interactions for the closed configuration were obtained from the MD simulations of the closed configuration model proposed in (13). We do not account for the electrostatic interaction between the negatively charged GLY and ASP residues as it remains unchanged during the S4 movement. Similarly, the electrostatic interaction between the ARG residues are not considered as there is no relative movement between them.

The salt bridge interaction energy is given by:

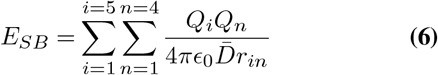

where 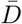 is the average dielectric constant for the salt-bridge forming charges, *Q_i_* are the positive charges, *Q_n_* are the counter charges, and *r_in_* is the separation between the saltbridge forming charges. The dielectric constant is assumed to vary across the bilayer as a hyperbolic tangent function ((Eqn. [S6]) as proposed in (65). As a result, the dielectric constant changes as the ARG residues move across the membrane.

#### ARG-anionic lipid interactions (*E_Lip_*)

We model the electrostatic energy between the clustered anionic lipids in the inner solvation shell and the positive VSD residues similar to the salt-bridge energy. The number of anionic lipids around the S4 and their distance from the positive residues are obtained from the RDF of anionic lipids computed from MD. This interaction energy is computed as

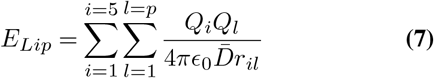

where p is the number of negatively charged lipid head groups around the protein, *Q_l_* is the charge of the lipid head group, *r_il_* is the separation between the charges, and 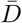 is the average dielectric constant for a positive residue-anionic lipid pair.

#### Self-energy of the charges (*E_self_*)

The self energy accounts for the energy required to move a charge from one dielectric medium to another. For a bilayer with varying dielectric constant (Eqn. [S6]), we use the model proposed in (65) to calculate the self-energy of the positive residues as

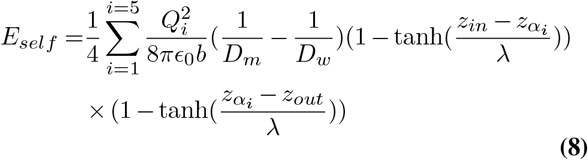

where *b* = 1.8 Å is the assumed radius for ARG and LYS residues, and *Q_i_* is the charge of the ARG/LYS residues.

### Mechanical Energy

The mechanical energy is associated with the bending and the thinning deformations of the membrane and is given by

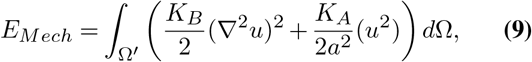

where the first term penalizes the bending deformations and the second term penalizes the thinning deformation of the membrane (20). Above, *K_B_* is the bending modulus, *K_A_* is the compression modulus, 2*u* is the change in membrane thickness, and 2*a* is the resting height of the membrane (see Figure S16; values listed in Table S5). Above, we have also assumed that the membrane is at zero resting tension. As shown in (20), employing the solution of the Euler-Lagrange equation (discussed in SI), we can express the mechanical energy as

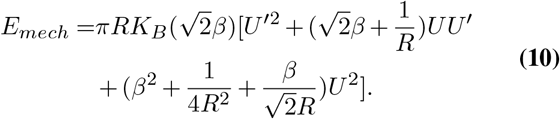

where *R* is the channel radius at the midplane, *U* is the thinning at the channel interface, *U*′ is the slope at the channel interface, and *β* = (*K_A_*/*K_B_a*^2^)^1/4^.

### Probability Distribution

The opening probability of the channel is calculated using a multi-state model given by

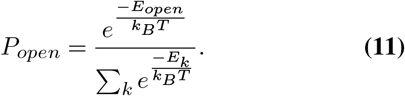

where *k* accounts for all the energy wells in the energy landscape, *E_k_* is the energy of a well and *E_open_* is the energy of the well corresponding to the open state, *k_B_* is the Boltzmann constant and *T* is the temperature.

## Acknowledgments

This work was supported by National Science Foundation Grants CMMI 1562043, CMMI 1727271 and CMMI 1931084 to A.A.. K.K.M was supported by Director, Office of Science, Office of Basic Energy Sciences, of the U.S. Department of Energy under contract No. DEAC02-05CH11231. The authors acknowledge the use of the Maxwell/Opuntia/Sabine Clusters and the advanced support from the Research Computing Data Core at UH to carry out the research presented here.

## Author contributions

A.A. conceived the study, N.T, K.K.M. and A.A. designed the MD simulations and the continuum analysis, N.T. carried out the atomistic and the continuum simulations, N.T, K.K.M. and A.A. analyzed the data and wrote the manuscript.

