## Supplemental Document for "Electromechanics of lipid-modulated gating of Kv channels"

### Molecular dynamics simulations

We used CHARMM-GUI (1, 2) to create all the systems. In addition to the description provided in the Methods sections, we outline additional details here. We added KCl salt to neutralize the simulation box. In the case of pure POPC system, we added two $Cl^-$ as POPC lipids are zwitterionic and protein has net two positive charges. Addition of 25% of CHOL resulted in a reduction of lipid diffusion. Therefore, temperature of POPC-CHOL system was increased to 323K in comparison to 303K for the rest of the systems. Simulations were performed using GROMACS 2018.3 (3). CHARMM36m (4, 5) force field was used throughout the simulations. Energy minimization of the system was performed using steepest descent method to bring the maximum force in the system down to less than 700 KJ/mol.nm. Energy minimization was followed by three sets of NVT and NPT equilibration cycles, respectively. Hydrogen bonds were constrained using LINCS algorithm (6). Verlet cut-off scheme was used with a cut-off distance of 1.2 nm. Short-range electrostatic interactions were cut-off at 1.2 nm and long-range interactions were calculated using PME (7). Van der Waal’s interactions were cut-off at 1.2 nm and a force-switch VdW modifier was used to reduce long-range VdW force to zero. Berendsen thermostat and barostat were used during the equilibration cycles with a time constant of 1.0 ps for temperature coupling and a time constant of 5.0 ps for pressure coupling. Semi-isotropic pressure coupling with different pressure coupling along z-direction was used throughout the simulation with compressibility of $4.5\times10^{-5}$ bar$^{-1}$. During NVT and NPT equilibration cycles, $C_\alpha$ carbons of VSD and phosphorus atoms in lipids were restrained and later the restraint forces were reduced to zero in production runs. The total NVT and NPT equilibration time for each system was 4.5 ns. The Nose-Hoover thermocouple (8, 9) was used in the production runs with a time constant of 1.0 ps. In addition, Parrinello-Rahman pressure couple (10) with 5.0 ps time constant and compressibility of $4.5\times10^{-5}$ bar$^{-1}$ was used to maintain the pressure at $10^5$ bar. The electrostatic interactions between the ARG residues and the negatively charged GLU and ASP residues were calculated using the VMD plugin ‘Salt Bridges’ (11). A cut-off distance of 0.35 nm used to define the donor-acceptor pair for a salt bridge (Figure S9). This data was also used to identify the significant salt-bridge connections between the S4 and S1-S2-S3 protein segments.

### The continuum model

#### Kinematics

The kinematics of the protein segments that regulates the gating of the ‘model’ channels comprises of two key contributions. First is the motion of the S4 segment in the VSD. It controls the motion of the ARG residues within the membrane and hence, controls the salt-bridge connections ARG residues make with the counter charges and the anionic lipids in the open and closed states. Second is the motion of the S6 segments in the PD. It controls the opening of the channel and the deformation of the membrane. These contributions are discussed below in greater detail.
**S4 motion:** The S4 segment is assumed to be a rigid cylinder representing the alpha-helix state of the protein and is oriented in the vertical direction. Figure S2 shows the side and the top views of the ‘model’ S4 in the closed and

open configurations for the KvAP channel. The radius of the S4 helix is considered to be 10 Å to account for the effective size created by the S4 side chains. The S4 geometry determines the coordinates of the ARG residues. The ARG residues in the S4 segment are distributed along the cylinder as individual charged beads (blue circles in Figure S2). We consider five ARG residues R117, R120, R123, R126 and R133 in the KvAP model. They are identified respectively as {3,2,1,0,-2.33} in the model. We suppress the interactions of the LYS located at K136 since it remains buried in the lower leaflet in both the open and the closed configurations.

We place the positive charges at the position of the $C_\alpha$ atoms of the ARG residues. The index of R133 is -2.33 as it is the seventh residue after R126. Each helical turn corresponds to 3.6 amino acids and the pitch of the alpha helix is assumed to be 5.4 Å. Therefore, the $C_\alpha$ atom of every residue moves by 4.5 Å down and rotates by 300 degrees in one pitch. We model the rotation of S4 about its axis as it undergoes vertical translation. The three dimensional coordinates of R117, R120, R123 and R126 residues are given by

$$x_{\alpha_i} = R_a \cos(\frac{\pi}{3} - \frac{\pi\alpha_i}{3} + \omega), \quad \text{[S1]}$$

$$y_{\alpha_i} = R_a \sin(\frac{\pi}{3} - \frac{\pi\alpha_i}{3} + \omega), \quad \text{and} \quad \text{[S2]}$$

$$z_{\alpha_i} = z + 4.5\alpha_i \quad \text{[S3]}$$

where $R_a = 10$ Å is the radius of alpha helix, $\alpha_i$ corresponds to the ARG index from the set {3,2,1,0,-2.33} for the KvAP channel, and $\omega$ and $z$ are the parameters used to define the rotation and the translation of the S4 helix. $z$ is defined as the position of R126 within the membrane and it varies from -9 Å to 9 Å to avoid complete exposure of peripheral ARGs into water. $\omega$ varies from 0 to 180 degrees. These assumptions are inspired from the model proposed by (12). For R133, the (x,y) coordinates are obtained by substituting $\alpha_5 = -1$ and the z coordinate is obtained by substituting $\alpha_5 = -2.33$. While the vertical position of R133 follows the normal alpha-helix rule, the (x,y) position of R133 is not in a standard place because of the kink in the S4 segment. Hence, the adjustment is required to obtain the correct coordinates. We would also like to note that the S4 helix axis is tilted by about 45° with respect to bilayer normal. However, we suppress the tilt in our model as ARG residue side chains are assumed to snorkel into the bilayer-water interface, which effectively diminish the effect of tilt.

The neighboring S1-S2-S3 domains contain four counter charges E107, E45, D62 and D72 with -1 charge. These include the GLU residues present on the S1 and S3 segments and the ASP residues present on the S2 segment. E107 is present on the S3b segment. The positive charges on the S4 and the negative charges on the S1-S2-S3 form salt bridge connections. Since S1 and S2 segments undergo minimal movement (13, 14, 15, 16, 17), we place these counter charges at fixed locations within the bilayer (red circles in Figure S2). The (x,y,z) coordinates of the counter charges are estimated based on the information of the salt-bridge interactions from (17).

**PD motion:** The S6 segments, which constitute the pore domain (PD), regulate the geometry of the lipids in the vicinity of the channel (Figure S3). We model the projected configuration of the S6 proteins. In the closed configuration, we assume that the S6 segments are oriented at 30° with respect to the vertical axis ($\theta_C^T = \theta_C^B = 30°$) as shown in Figure S3a. In the open configuration, the protein segments in the inner leaflet (C-terminal segment) are assumed to rotate to 45° ($\theta_O^B = 45°$) (Figure S3b). These assumptions are qualitatively based on the findings presented in (18, 13, 15, 19, 20, 21). Based on the hypothesized model proposed in (22), we assume that the protein segments in the outer leaflet (N-terminal segment) maintains the same orientation in both the open and the closed configurations. Looking from the top (Figure S3c and Figure S3d), each S6 segment is assumed to be oriented at 15°($\phi_C^T = \phi_C^B = 15°$) with respect to the radial vectors connecting the pore center to the N-terminus of the S6 segment. In the open configuration, the C-terminal segment is assumed to be twisted to 45°($\phi_O^B = 45°$) but the twist of the N-terminal segment is assumed to remain unchanged (based on (22)). We approximate the PD comprising of discrete helices with a hollow cone (Figure S3e and S3f). In the open configuration, the cone has a kink at the midplane because of the different tilt angles of the S6 segments in the outer and inner leaflets. We would like to note that the protein-membrane interface becomes slightly curved in the open configuration due to tilting and twisting of the protein segments (see Figure S3d). But we approximate the interface with straight edges in the model.

The motion of the S4 and the S6 segments are linked during the gating of the channel. We therefore link the instantaneous tilt and twist angle associated with the S6 segment to the S4 kinematics through the mappings

$$\theta^B = \theta_C^B + \frac{\theta_O^B - \theta_C^B}{z_O - z_C}(z - z_C), \quad \text{[S4]}$$

and

$$\phi^B = \phi^B_C + \frac{\phi^B_O - \phi^B_C}{\omega_O - \omega_C}(\omega - \omega_C), \quad \text{[S5]}$$

where we used $\theta^B_C = 30°$, $\theta^B_O = 45°$, $z_C = -9$Å $z_O = 9$Å, $\phi^B_C = 15°$, $\phi^B_C = 45°$, $\omega_C = \pi$, and $\omega_O = 0$ for the KvAP channel.

#### Dielectric constant

The dielectric constant variation was modeled by (12)

$$D = D_w + (\frac{1}{4}(D_m - D_w)(1 - \tanh(\frac{z_{in} - z}{\lambda}))(1 - \tanh(\frac{z - z_{out}}{\lambda}))), \quad \text{[S6]}$$

where $D_w = 80$ is the dielectric constant of water, $D_m = 14.5$ is the dielectric constant of lipid membrane assumed in the vicinity of KvAP channel, $\lambda$ is a length constant that regulates the rate of change of $D$, $\{z_{in}, z_{out}\}$ are the z-coordinates of the water interfaces inside the membrane, and $z$ is the position of the counter charge or the positive residue determined from Eqn. [S3]. The dielectric constant plot is shown in Figure S11.

#### Mechanical Energy

The mechanical energetic cost is given by

$$E_{Mech} = \int_{\Omega'} \left( \frac{K_B}{2}(\nabla^2 u)^2 + \frac{K_A}{2a^2}(u^2) \right) d\Omega, \quad \text{[S7]}$$

where the first term penalizes bending deformations and the second term penalizes membrane thinning. Above, $K_B$ is the bending modulus, $K_A$ is the compression modulus, $2u$ is the change in membrane thickness, and $2a$ is the resting height of the membrane.

The variation of Eqn. [S7] with respect to $u$ yields the equilibrium equation

$$K_B \nabla^4 u + \frac{K_A}{a^2} u = 0. \quad \text{[S8]}$$

Above, we have suppressed the energetic contribution due to the bending of the membrane midplane as it is negligible (also shown to be negligible in (23)). Since we require the system to possess cylindrical symmetry, the equilibrium solution is given by the zeroth order modified Bessel functions of the second kind (23).

$$u(r) = A_+ K_0(\beta_+ r) + A_- K_0(\beta_- r) \quad \text{[S9]}$$

where

$$A_\pm = -\frac{K_\mp U' + \beta_\mp U K'_\mp}{\beta_\pm K'_\pm K_\mp - \beta_\mp K'_\mp K_\pm}, \quad \text{[S10]}$$

$r$ is the radial distance, $2U$ is the thinning of membrane at the membrane-protein interface ($U = u(R)$), and $U'$ is the slope of the membrane-protein interface ($U' = \frac{1}{2}(\theta^T - \theta^B)$ in the open and closed states, $\beta_\pm = e^{\pm\frac{\pi i}{4}}\beta$, $\beta = (K_A/K_B a^2)^{1/4}$, $K_\pm = K_0(\beta_\pm R)$ and $K'_\pm = K'_0(\beta_\pm R)$.

As shown in (23), the total energy in Eqn. [S7] can be reduced to the boundary integral

$$E_{mech} = \frac{1}{2} \oint_{\partial\Omega'} d\hat{n}.(K_B[\nabla u \nabla^2 u - u \nabla^3 u]. \quad \text{[S11]}$$

Substituting Eqn. [S9] into Eqn. [S11] we get

$$E_{Mech} = \pi R K_B \frac{(\beta_+^2 - \beta_-^2)(K_+ U' - \beta_+ U K'_+)(K_- U' - \beta_- U K'_-)}{\beta_- K_+ K'_- - \beta_+ K_- K'_+}, \quad \text{[S12]}$$

where R is the radius of protein at the midplane. Substituting the expressions for $\beta_{\pm}$, $K_{\pm}$ and $K'_{\pm}$, we get

$$E_{mech} = \pi R K_B(\sqrt{2}\beta)[U'^2 + (\sqrt{2}\beta + \frac{1}{R})UU' + (\beta^2 + \frac{1}{4R^2} + \frac{\beta}{\sqrt{2}R})U^2]. \quad \text{[S13]}$$

For our calculations, we have considered a bilayer thickness of 4 nm and the projected length of the C-terminal and N-terminal of S6 segments to be 2 nm each in the closed state of the channel. Since the projected length of the S6 segments is equal to the leaflet thicknesses, there is no membrane thinning in the closed configuration and $U = 0$. The headgroup-tail interface of each leaflet is assumed to be perpendicular to the projected S6 segments (see Figure S10). In the closed configuration, the S6 segments are tilted 30° with respect to vertical axis ($\theta_C^T = \theta_C^B = 30°$ in Figure S3c). Since the slope of both the outer and the inner leaflets are identical at the channel interface, $U' = 0$ in the closed configuration. The twist angle of the S6 segments is assumed to be 15° with respect to the radial vector connecting the center of the PD and the N-terminal end of the S6 segment ($\phi_C^T = \phi_C^B = 15°$ in Figure S3e). In the open configuration, the C-terminal segment of S6 tilts to 45° ($\theta_O^B = 45°$ in Figure S3d) and twists to 45° ($\phi_O^B = 45°$ in Figure S3f). This results in the thinning of the inner leaflet $U = -1.835$ Å and a relative change in the slope of the lipids at the channel interface results in $U' = 0.124$. The membrane deformation resulting from the movement of the PD is shown in Figure S10.

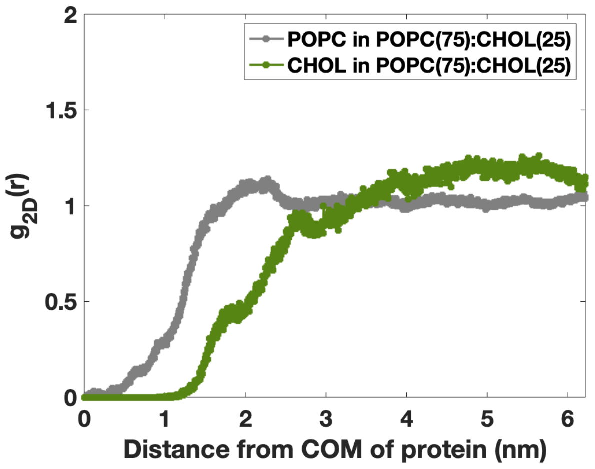


Figure S1: RDFs of POPC lipids (gray curve) and CHOL (gray curve) with respect to the COM of VSD in the outer leaflet of the membrane. CHOL does not show aggregation around VSD in the outer leaflet.

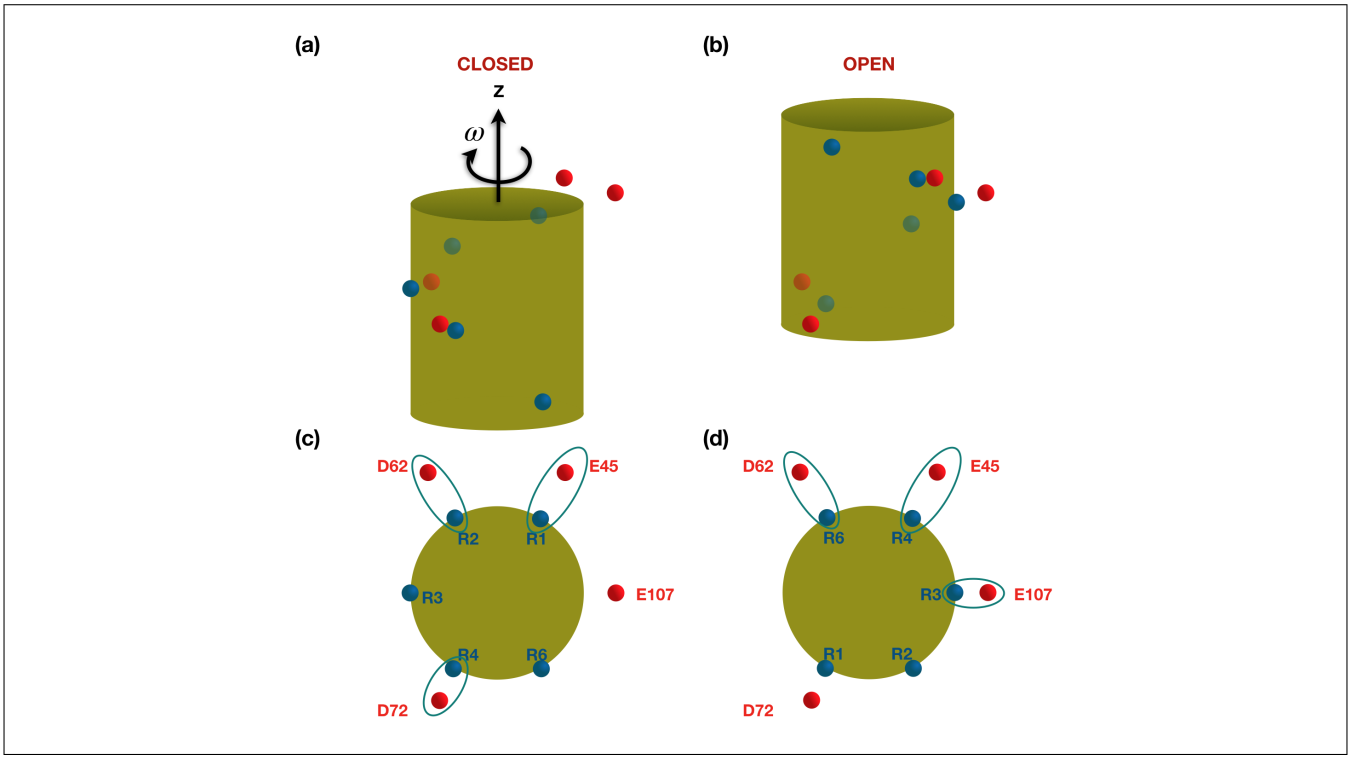


Figure S2: Kinematics and charge distribution of the 'model' KvAP S4 segment. (**a** and **b**) Front views of the S4 helix represented as a cylinder (in green) in the closed and open configurations, respectively. The ARG residues on S4 are shown as blue spheres and the counter charges are shown as red spheres. The S4 helix rotates by $\pi$ radians and moves upwards within the membrane during the transition from the closed state to the open state. (**c** and **d**) Top views corresponding to (**a**) and (**b**) showing the salt bridges connections in the closed and open states (blue ellipses).

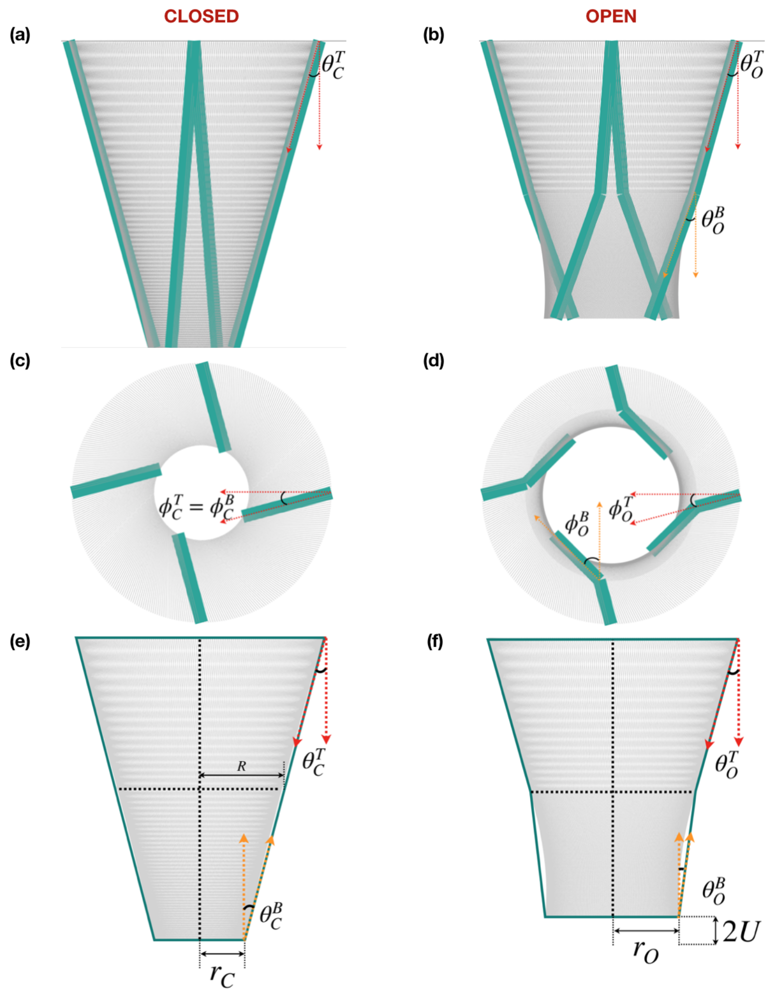


Figure S3: Kinematics of the ‘model’ PD. (**a** and **b**) Orientation of four S6 segments in the closed and open configurations (side view). In the closed configuration, S6 segments are assumed to be tilted at 30 °with respect to the vertical axis ($\theta_C^T = \theta_C^B = 30°$). In the open configuration, the C-terminal segment of the S6 segment in the inner leaflet is assumed to tilt to 45° ($\theta_O^B = 45°$). The N-terminal segment of the S6 segment in the outer leaflet is assumed to maintain the same tilt. (**c** and **d**) Top views of the four S6 segments in the closed and open configurations. Each segment is assumed to be oriented at 15° ($\phi_C^T = \phi_C^B = 15°$) with respect to the radial vectors connecting the pore center to the N-terminus of the segment. In the open configuration, the N-terminal segment of the S6 is assumed to remain at ($\phi_O^T = 15°$) but the C-terminal segment is assumed to twist to 45°($\phi_O^B = 45°$). These assumptions are qualitatively based on the findings presented in (22, 19). (**e** and **f**) The PD is modeled as (solid) hollow conical sections in the outer and the inner leaflets. The angles of the conical interfaces are obtained from the orientations of the S6 protein segments. In the open configuration, the PD has a kink at the midplane because of the different tilts of the S6 segments in the two leaflets. The open configuration also results in the thinning of the inner leaflet.

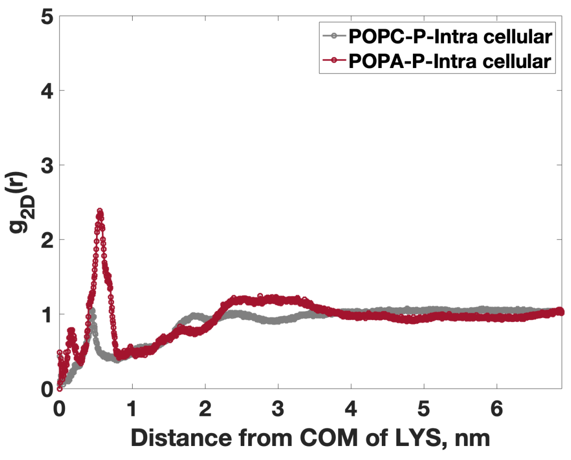


Figure S4: RDFs of POPA lipids (red curve) and POPC lipids (gray curve) with respect to the COM of LYS136 residue in the inner leaflet of the membrane. The peak in the red curve demonstrates preferential aggregation of POPA lipid around LYS in the inner leaflet even in the open configuration of the VSD.

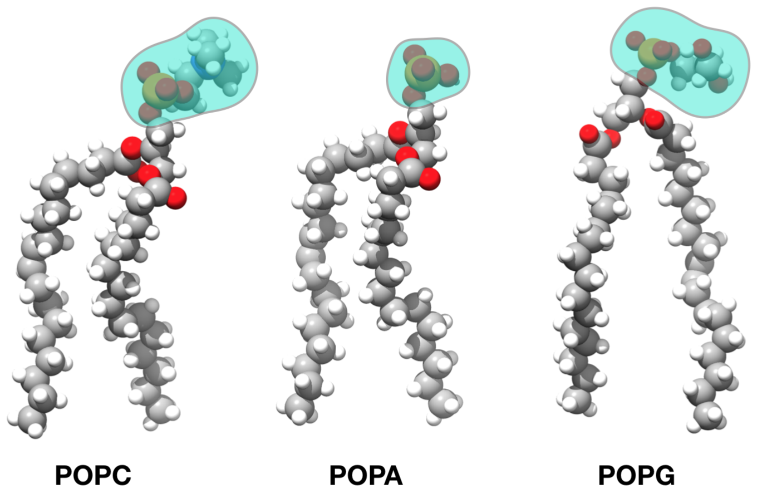


Figure S5: Comparison of the headgroup sizes of POPC, POPA, and POPG lipids. Volumetric footprint of the POPA headgroup is 35% smaller than those of the POPC or POPG lipids. This leads to a higher charge density in the POPA headgroup compared to the POPG headgroup, which consequently leads to stronger electrostatic interactions with the ARG residues of the S4 segment.

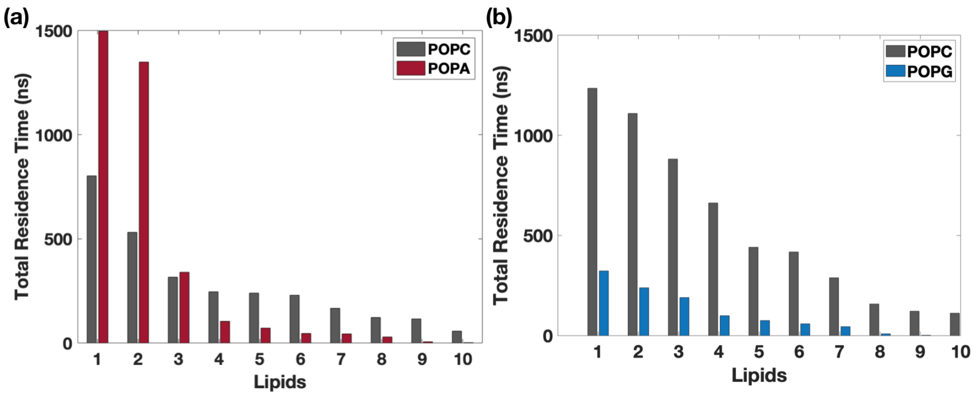


Figure S6: POPA lipids spend more time in the vicinity of S4 than POPG lipids (**a**) Total residence time of ten POPC and POPA lipids that show strong interactions with the ARG residues in the mixed POPC(75):POPA(25) membrane. (**b**) Total residence time of ten POPC and POPG lipids that show strong interactions with the ARG residues in the mixed POPC(75):POPG(25) membrane. The time POPA lipids spend is significantly larger than the time POPG lipids spend in the vicinity of S4, indicating that POPA lipids exhibit stronger direct electrostatic interactions with ARG residues than POPG lipids. Since both POPA and POPG lipids have the same unit negative charge and acyl chains, POPG's reduced electrostatic interactions with the ARGs could be attributed to its bigger headgroup size. The total residence time is computed as the total time a lipid phosphorus atom spends within 5 Å of any atoms of R117, R120, R123 and R126 residues.

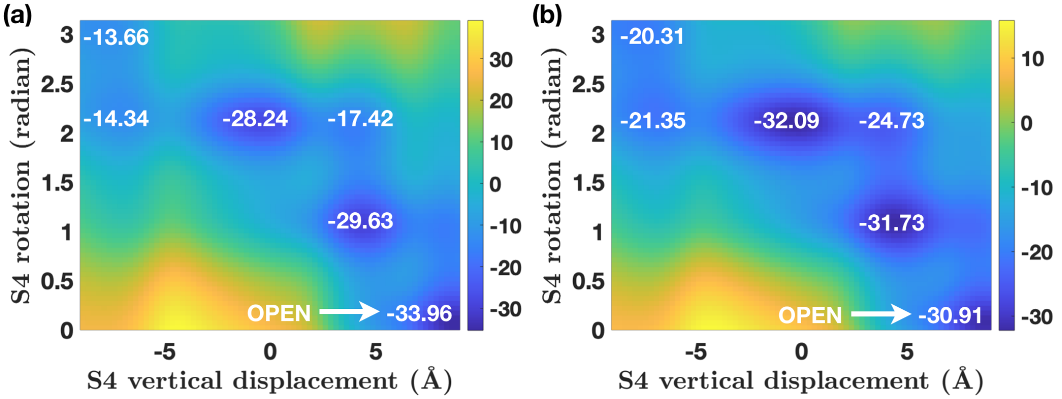


Figure S7: Predicted effect of DOTAP on KvAP gating. (**a** and **b**) Energy landscapes of the KvAP channel at 0 mV transmembrane potential in pure POPC and mixed POPC-DOTAP membranes. The vertical position of S4 is on the x-axis, the rotation of S4 is on the y-axis, and the energy values (in $k_B T$) at the wells are in white text. We increased the dielectric constant to simulate the effect of DOTAP lipids. This is based on the finding that DOTAP desolvates the S4 triggering higher water penetration in the protein vicinity (24). As a result, electrostatic interactions diminish and higher depolarization is needed to open the channel.

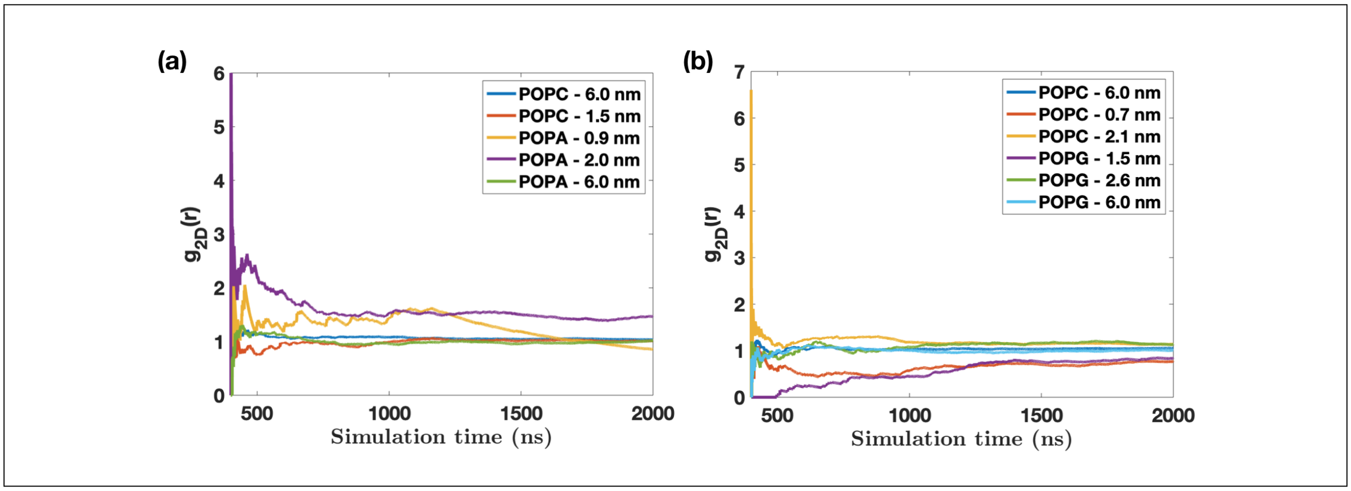


Figure S8: RDFs as a function of time in the outer leaflet evaluated at different radial distances from the COM of VSD for (**a**) POPC(75):POPA(25) and **b** POPC(75):POPG(25) systems. By the end of simulations, the fluctuations fall below 10%, showing convergence of RDFs.

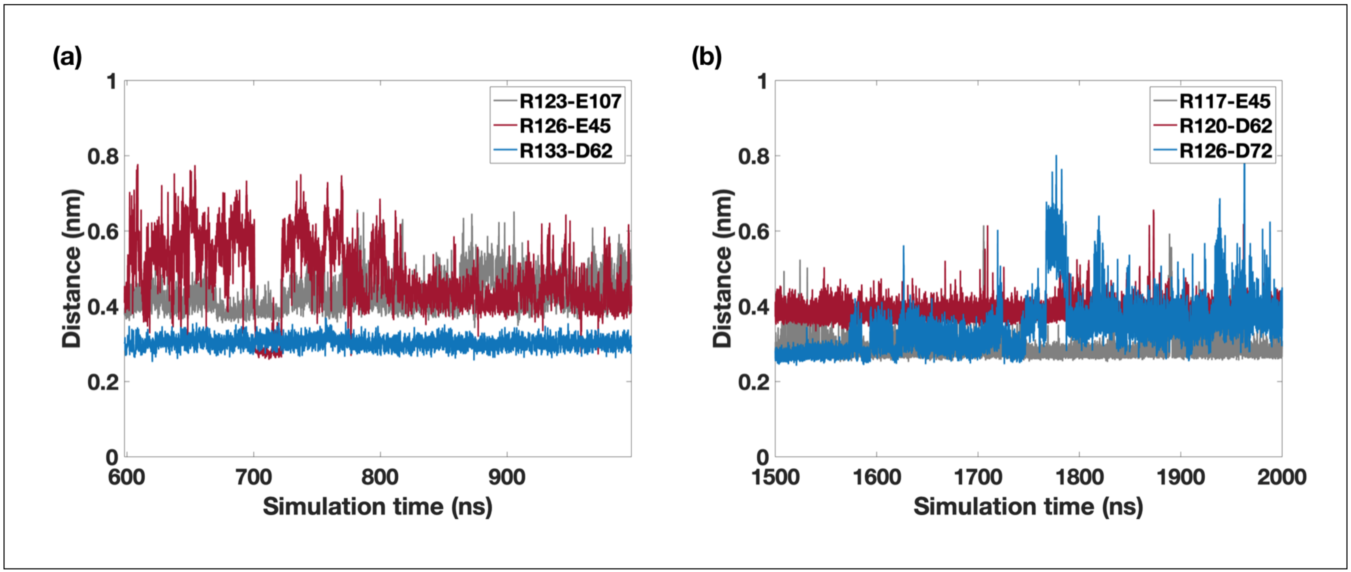


Figure S9: Salt bridge analysis for the KvAP channel in the open (**a**) and closed (**b**) configurations. Distances are measured from the COM of nitrogen atoms of ARG residues and oxygen atoms of ASP and GLU residues. The analysis reveals three stable salt bridges in both the configurations.

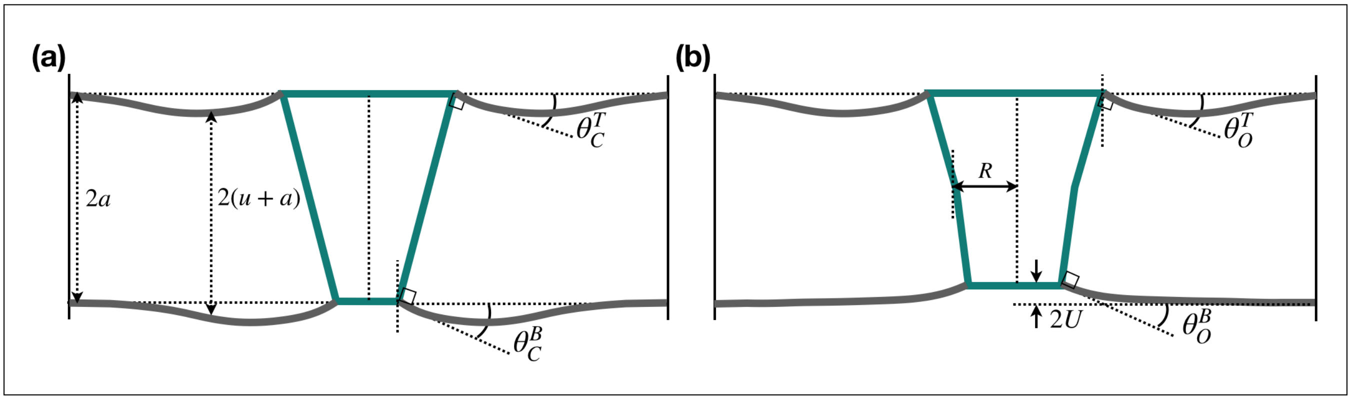


Figure S10: Membrane deformation in the closed and open configurations of the 'model' Kv channel. (**a**) In the closed configuration, we assume that that projected height of the conical PD is equal to the resting thickness of the membrane ($2a = 4$ nm). As a result, $U = 0$. Since the tilt of the conical sections in the outer and inner leaflets are equal, $U' = \frac{1}{2}(\theta_C^T - \theta_C^B) = 0$ in the closed configuration. (**b**) In the open configuration, $U = -1.835$ Å and $U' = 0.124$ as the conical section in the inner leaflet tilts to $\theta_O^B = 45°$ and twists to $\phi_O^B = 45°$.

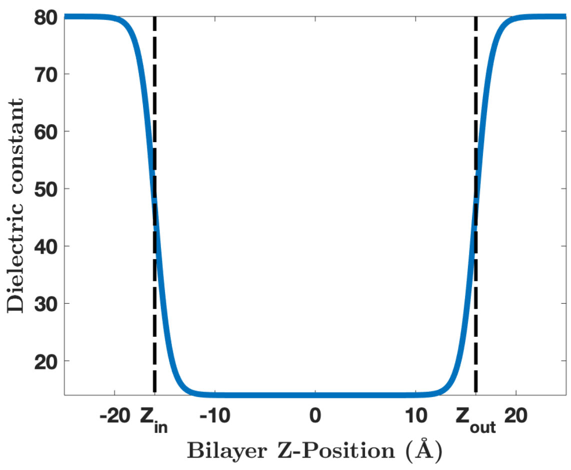


Figure S11: Dielectric constant variation across the lipid bilayer. $z_{in}$ and $z_{out}$ are the z-coordinates of the water interfaces inside the membrane and are marked as dotted lines. The dielectric constant of lipid bilayer in the vicinity of proteins is assumed to be 14.5 and the dielectric constant for water is 80.

Table S1: Summary of salt bridge interactions for KvAP channel

| **Salt bridge** | **State of channel** | **Distance between charges (Å)** |
|---|---|---|
| E107-R123 | Open | 3.64 |
| E45-R126 | Open | 6.0 |
| D62-R133 | Open | 6.0 |
| E45-R117 | Close | 6.0 |
| D62-R120 | Close | 7.21 |
| D72-R126 | Close | 4.0 |

Table S2: Summary of coordinates of counter charges and anionic lipids used in the continuum analysis of KvAP channel

| **KvAP channel parameters** | |
|---|---|
| Coordinates of E45 | (8,13.59,12.5)Å |
| Coordinates of D62 | (-9,14,-4)Å |
| Coordinates of D72 | (-9,-15.0,-7)Å |
| Coordinates of E107 | (13.8,0,13)Å |
| Coordinates of $1^{st}$ PA lipid in outer leaflet | (-10.72,-9.00,20)Å |
| Coordinates of $2^{nd}$ PA lipid in outer leaflet | (-3.47,-19.70,20)Å |
| Coordinates of $1^{st}$ PA lipid in inner leaflet | (2.48,-13.77,-19)Å |
| Coordinates of $2^{nd}$ PA lipid in inner leaflet | (9.92,-8.39,-19)Å |
| Coordinates of $3^{rd}$ PA lipid in inner leaflet | (10.0,-17.32,-19)Å |

Table S3: Summary of parameters used in the continuum model of KvAP channel

| **Parameter** | **Values** | **Source** |
|---|---|---|
| Dielectric constant of water ($D_w$) | 80 | (25) |
| Dielectric constant of lipids-KvAP ($D_l$) | 14.5 | (26) (27) |
| Radius of S4 ($R_a$) | 10 Å | |
| Length constant ($\lambda$) | 1.5 Å | |
| Inner leaflet interface ($Z_{in}$) | -16 Å | |
| Outer leaflet interface ($Z_{out}$) | 16 Å | |
| Radius of positive charged residue (b) | 1.8 Å | |
| Monolayer thickness (a) | 20 Å | (28) |
| $K_B$ for POPC system | 27 $k_BT$ | (29) (28) |
| $K_B$ for POPC:CHOL system | 50 $k_BT$ | |
| $K_A$ for POPC system | 0.620 $k_BT/Å^2$ | (29) (28) |
| $K_A$ for POPC:CHOL system | 1.569 $k_BT/Å^2$ | (30) |

Table S4: Summary of the CRAC/CARC domains in the KvAP channel

| **Domain** | **Segment** | **Leaflet** | **1$^{st}$ Residue** | **2$^{nd}$ Residue** | **3$^{rd}$ Residue** |
|---|---|---|---|---|---|
| CARC | S5 | in | LYS(147) | TYR(151) | VAL(157) |
| CARC | S5 | in | ARG(149) | PHE(154) | LEU(159) |
| CARC | S5-S6 | out | LYS(181) | PHE(184) | LEU(187) |
| CRAC | S6 | in | VAL(231) | PHE(234) | LYS(237) |

Table S5: Summary of the molecular dynamics simulations performed in this study

| **Bilayer composition** | **Lipids per leaf** | **Simulation time ($\mu$s)** |
|---|---|---|
| POPC-Pure | POPC (200) | 1.0 |
| POPC-POPA | POPC (225): POPA (75) | 2.0 |
| POPC-POPG | POPC (225): POPG (75) | 2.0 |
| POPC-CHOL | POPC (226): CHOL (75) | 2.0 |
| POPC-CHOL-Mut | POPC (226) : CHOL (75) | 1.4 |
